# Decoding CBM2 binding modes: how similar folds achieve distinct substrate preferences

**DOI:** 10.64898/2026.10.13.758739

**Authors:** Susanne Hansen Troøyen, Andrew Philip Rennison, Tina Dahlgren, Peter Fojan, Marie Sofie Møller, Gaston Courtade

**Affiliations:** Norwegian University of Science and Technology (NTNU); Technical University of Denmark (DTU); Aalborg University (AAU)

## Abstract

Understanding the mechanisms underlying substrate recognition and selectivity is essential for developing new biotechnological applications of carbohydrate-binding modules (CBMs). Many members of CBM family 2 (CBM2) prefer binding to crystalline substrates and show diverse substrate specificities despite sharing similar folds. We show here that CBM2 specificity cannot be rationalized solely based on sequence and fold. Rather, specific binding of CBM2s to crystalline surfaces depends on a balance of topological complementarity, hydrophobic dehydration, and hydrogen-bonding interactions. We compare the binding modes of *Sc*CBM2 and *Ba*CBM2 on cellulose, chitin, and polyethylene terephthalate (PET) with respect to protein and substrate properties. Using MD simulations and NMR, we explore the CBMs’ surface topology, hydration, solvent accessibility, and electrostatics. The results indicate that the two CBMs may bind to crystalline substrates in opposite orientations, and that the interplay between hydrophobic and π-stacking interactions along the aromatic triad and hydrogen-bonding interactions on either side determines their orientation. We further show that higher water density at the aromatic triad correlates with higher substrate affinity. Lastly, we probe the interaction dynamics of *Ba*CBM2 with PET nanoparticles and the monomer bis(2-hydroxyethyl) terephthalate (BHET) using NMR relaxation and solvent paramagnetic relaxation enhancement (sPRE) experiments, revealing a mixture of transient, low-population states involved in PET binding. These insights clarify the interplay among factors that drive CBM2 substrate recognition and highlight the importance of evaluating surface hydration and protein dynamics in future CBM2 studies.

## Introduction

Carbohydrate-binding modules (CBMs) are non-catalytic protein domains commonly appended to carbohydrate-active enzymes (CAZymes) [1]. They enhance the catalytic activity of their associated catalytic domains through substrate recognition and binding, promoting prolonged contact with their carbohydrate substrates [2, 3]. CBM family 2 (www.cazy.org) is a large family of CBMs, primarily found in bacterial CAZymes, that have been shown to bind cellulose, chitin, or xylan [4, 5]. Within this family, there are type A CBMs that bind with high affinity to crystalline substrates due to their characteristic planar binding surface, and type B CBMs harboring a binding groove that enables *endo* binding to individual glycan chains [1]. Two antiparallel β-strands arranged in a β-jelly-roll fold comprise the canonical CBM2 structure, and, in type A CBM2s, three highly conserved aromatic residues (the aromatic triad) make up the planar binding site [1, 5].

Exploring CBM binding modes is complicated by substantial structural variation among substrates. Cellulose is the most abundant natural polysaccharide on Earth and can adopt diverse microstructures, crystallinities, and crystal morphologies. CBMs have evolved to target these structural variations with varying degrees of selectivity, some being highly specific and others promiscuous [6–10]. Even within a single cellulose sample, several cellulose forms may coexist, making the characterization of CBM binding complicated. Typically, there is not a uniform distribution of equally accessible binding sites, but rather a mix of binding sites to which different CBMs display different binding modes and specificities [11, 12].

Several CBMs have been shown to bind the synthetic polymer polyethylene terephthalate (PET), underscoring the diversity of CBM selectivity [13, 14]. This promiscuity has been exploited by fusing CBMs to PET-hydrolyzing enzymes to enhance their activity [15–21]. However, some studies show that CBMs have negligible impact at high substrate loads relevant to industrial recycling processes [17]. Still, CBMs can bind a wide range of substrates, raising the question of what mechanisms enable recognition of both natural and synthetic polymer surfaces.

CBM binding to a crystalline polymer surface requires dehydration to create sufficient contact for π-stacking and hydrogen bonding [11, 22]. This removal of structured water is associated with positive entropy changes because it increases the degrees of freedom of the water molecules. For soluble substrates, the favorable entropy contribution of dehydration is outweighed by a decrease in entropy arising from conformational restriction of the substrate and protein in the bound state. Hence, CBM-oligosaccharide interactions tend to be driven by mechanisms that induce negative enthalpy changes, such as π-stacking and hydrogen bonding [23]. For insoluble substrates, the substrate conformation is already restricted, allowing the positive entropy change from dehydration to drive binding. Creagh *et al.* showed through isothermal titration calorimetry (ITC) that binding of CBM2 from *Cellulomonas fimi* to bacterial microcrystalline cellulose was entropically driven, and that dehydration at the CBM-cellulose interface was the dominant contributor to the increased entropy upon binding [11]. A similar entropy effect was demonstrated by Zhang *et al.* for CBM1 and CBM3 domains binding to cellulose nanofibrils (CNF) and cellulose nanocrystals (CNC) [24].

However, the desolvation mechanism is not universal for all CBMs that bind to insoluble substrates. For example, binding of CBM17 from *Clostridium cellulovorans* and CBM28 from *Bacillus* species 1139 to amorphous cellulose, and CBM1 from *Trichoderma reesei* to crystalline cellulose have been shown to be largely driven by decreased enthalpy [8, 25]. Liu *et al.* demonstrated that CBM4 from *Cellulomonas fimi* binding to CNF was mainly driven by enthalpy, whereas binding to CNC was dominated by favorable changes in entropy. They attributed the different binding modes to lower amorphous content in the CNC and higher amorphous content in the CNF, leaving more cellulose chains available for *endo* binding in the CNC material [26].

Because CBMs adopt highly specialized binding specificities, their potential applications in biotechnology are vast. CBM applications include enzyme immobilization, CBM-enzyme fusion design, and functionalized materials design [3, 27, 28]. Developing these applications requires a deeper understanding of how structurally related CBMs within the same family achieve distinct substrate specificities across both natural polysaccharides and synthetic polymers, a relationship that remains poorly understood.

Recently, a high-throughput assay for screening CBM binding selectivity was established and applied to screen the binding of 800 CBMs from selected families, including CBM2, to cellulose, chitin, starch, PET, polystyrene (PS), and polyethylene (PE) [21]. Screening large numbers of binding modules provides invaluable insight into how specificity or promiscuity correlates with phylogeny and structure, as discussed in their work. To complement these large studies, we need comparative studies that explore the binding modes of selected CBMs. Our study builds on this by offering a complementary, in-depth characterization of two type A CBM2s with distinct substrate specificities. To investigate how the same CBM fold achieves differential substrate selectivity, we selected *Sc*CBM2 of the cellulose-active lytic polysaccharide monooxygenase *Sc*LPMO10C from *Streptomyces coelicolor* (GenBank: CAB61600.1) and *Ba*CBM2 of chitinase B from *Bacillus anthracis* (GenBank:

ACQ50287.1). We evaluate their differential binding to cellulose, chitin, and PET. For *Sc*CBM2, we include a more detailed inspection of cellulose binding, using cellulose substrates of varying crystallinity. To further explore factors governing selectivity, we present molecular dynamics (MD) simulations and NMR results, which reveal differences in water density and solvent accessibility at the CBMs’ binding sites. Finally, we probe residue-specific substrate interactions by using NMR to map chemical shift perturbations and nuclear relaxation effects. Together, these results advance our understanding of CBM2 binding mechanisms and how they likely depend on a complex balance of structure, surface hydration, and protein dynamics.

## Results and discussion

### Impact of amino acid sequence on substrate targeting in family 2 CBMs

To test whether substrate promiscuity in CBM2 is encoded at the sequence level, we performed a multiple-sequence alignment [29] of domains for which promiscuous or highly specific binding to cellulose, chitin, and PET has been experimentally characterized [21]. ***Figure S1*** shows sequence logos [30] of CBM2 modules that bind each substrate. The consensus sequences show no obvious differences among the three substrates, most likely because most CBM2s in the dataset are promiscuous binders and therefore appear in two or all three alignments. We observed the same consensus among CBM2s that bind both cellulose and PET (***Figure S2***).

Focusing on highly specific CBM2s that were confirmed to bind only a single substrate, we found that the residue at the third position of the aromatic triad more often deviates from the typical tryptophan residue (***Figure S3***). In cellulose- and chitin-specific CBM2s, this position is occupied by either tryptophan, tyrosine, or, in a few cases, histidine. This suggests that variation at the third position is particularly important for substrate specificity in CBM2s, which is consistent with the idea that tuning the aromatic triad is an efficient way to adapt the binding surface to a given polymer. However, residue identity does not determine whether the CBM is specific to cellulose versus chitin. We could not find any specific positions of interest that distinguish cellulose- and chitin-specific CBM2s; in fact, their sequence logos are strikingly similar, showing that specificity cannot be determined from sequence conservation alone.

The dataset of PET-specific CBM2s contained only six proteins, with very low conservation (***Figure S3***). In three of them, glutamine or asparagine occupies the third position of the aromatic triad, indicating a preference for polar residues at this site. Despite the large sequence variation, charged and polar residues appear more prevalent throughout PET-specific CBM2 sequences than in consensus sequences for promiscuous CBM2s. It has been shown that strong PET binding is correlated with polar patches separating the hydrophobic aromatic triad [21]. These observations suggest that hydrophilic residues capable of strong hydrogen bonding or electrostatic interactions may be important for specific PET targeting. Still, a high content of these residues is not a prerequisite for PET-binding ability in promiscuous CBM2s (***Figure S1***).

### Comparison of *Sc*CBM2 and *Ba*CBM2 fold and sequence

To establish a baseline for comparing *Sc*CBM2 and *Ba*CBM2 binding modes on polymer substrates, we first examined their fold and sequence features. Both CBMs adopt the characteristic β-jelly-roll fold (***Figure S4***-A), with the aromatic triad W13, W50 and H69 on *Sc*CBM2 (***Figure 1***-A), and W9, W44, W63 on *Ba*CBM2 (***Figure 1***-B) arranged on a planar strip. The sequence alignment (***Figure 1***-C) reveals 28 fully conserved residues, including two of the tryptophans in the aromatic triads. A key difference between the CBMs is the identity of the third aromatic residue: in *Sc*CBM2 it is histidine, and in *Ba*CBM2 it is tryptophan. We discuss the impact of an H69W substitution on *Sc*CBM2 binding to PET later.

**Figure 1.**
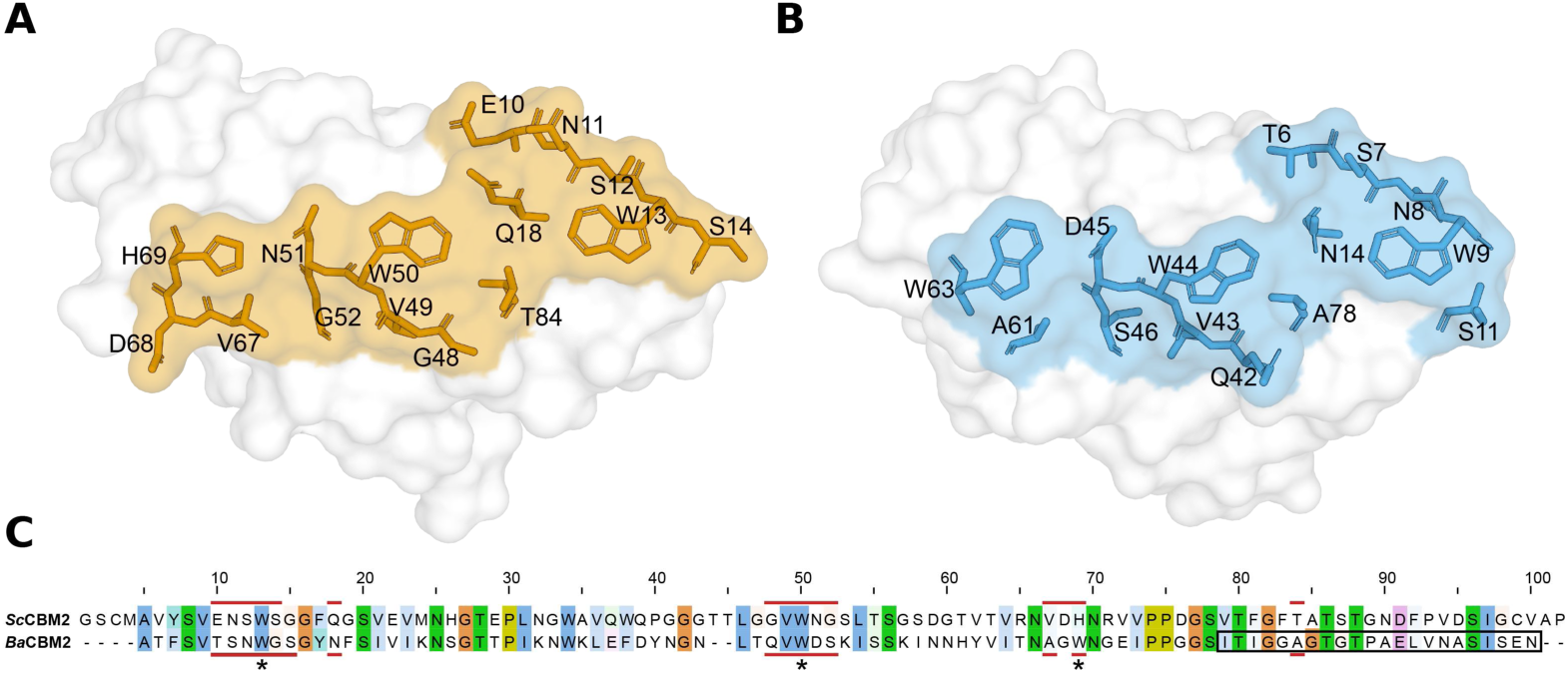
AlphaFold [31] structures of *Sc*CBM2 (A) and *Ba*CBM2 (B) with residues close to the planar strip harboring the aromatic triad (W13, W50, and H69 on *Sc*CBM2, and W9, W44, and W63 on *Ba*CBM2) shown as sticks. (C) Sequence alignment with conserved residues (60% identity) highlighted [29, 32]. Residues on the binding strip are marked with red lines to show their position on the sequences. The aromatic triad residues are marked with an asterisk (*). The 22 residues missing from the C-terminal truncated version *Ba*CBM2-t are marked with a black outline in the sequence.

Previous studies have used two versions of *Ba*CBM2 [13, 14]. In earlier work by Weber *et al.*, the PET-binding ability of a C-terminal truncated version (see ***Figure 1***-C), referred to as *Ba*CBM2-t in our work, was investigated [13]. In our binding studies, we used the full-length version, which has increased affinity, but lower binding capacity, to PET [14]. In the truncated version, *Ba*CBM2-t, one of the β-strands below the binding surface is removed, resulting in a more open and irregular binding site (***Figure S4***-B/C).

Using AlphaFold [31] structures, we compared the amino acid environments around the aromatic triad to identify distinguishing features of the putative binding surfaces. Among the nearest neighboring residues (***Figure 1***), only one residue, V49 in *Sc*CBM2 and V43 in *Ba*CBM2, is conserved, aside from the aromatic residues themselves. Several peripheral positions around the aromatic triad are occupied by residues with different properties in the two CBMs (***Figure S4***-C). Notably, the residues flanking W13 in *Sc*CBM2 (E10, S14, and T84) are more polar than those surrounding W9 in *Ba*CBM2 (T6, G10, and A78). Three residues on the β-sheet that carries the first two residues of the aromatic triad are significantly bulkier in *Sc*CBM2 (E10, N11, and Q18) than in *Ba*CBM2 (T6, S7, and N14), which may sterically restrict substrate binding. Between the second and third aromatic triad residues, *Sc*CBM2 harbors an uncharged asparagine (N51), whereas *Ba*CBM2 has a negatively charged aspartic acid (D45) that increases the polarity of the region between the aromatic residues. The extent to which these differences contribute to fine-tuning substrate selectivity remains unclear.

### Cellulose crystallinity and morphology

CBM binding to cellulose is sensitive to substrate crystallinity, crystal polymorph, and particle morphology. To interpret the binding measurements described below, we characterized the crystallinity and morphology of microcrystalline cellulose (Avicel), nanocrystalline cellulose (CNC), and cellulose nanofibrils (CNF) used in this study.

XRD patterns of the three types of cellulose are shown in ***Figure S5***. Earlier reports of cellulose crystallinity have often been based on the Segal method (see ***Figure S6*** for its application to our data), in which crystallinity is calculated from XRD peak intensities at only two 2θ angles, one amorphous and one crystalline [33]. However, this method has been widely disputed, partly because relying solely on peak intensities fails to account for peak area [34–36]. In cellulose, a small crystallite size usually yields broad XRD peaks, making it extremely important to consider peak area when determining crystallinity. To fit the XRD patterns, we therefore used a Rietveld method based on previously reported cellulose Iβ and II crystal structures [37, 38], and determined the amorphous fraction by including two peaks fixed at 20.52° and 38.87° 2θ, as explained by Ju *et al.* [39]. ***Table 1*** shows the calculated crystallinity index (CI) and crystallite size from our measurements.

**Table 1.** Cellulose crystallinity and crystallite size of cellulose substrates, determined by XRD using the Rietveld method. The cellulose polymorph is indicated in parentheses.

|  | Avicel | CNC | CNF |
| --- | --- | --- | --- |
| Crystallinity index (CI, %) | 85 | 91 | 46 |
| Crystallite size (nm) | 5.10 (Iβ) | 3.29 (Iβ), 14.7 (II) | 4.07 (Iβ) |

The most crystalline substrate was CNC (CI = 91%), in which we identified two distinct cellulose polymorphs. Peak areas after XRD data refinement showed that 65% of the total crystalline fraction is cellulose Iβ, and 35% is cellulose II. Cellulose II is formed during CNC production, typically following alkaline treatment after acid hydrolysis. AFM analysis (***Figure S7***) showed the CNC as disc-like particles of about 0.1-0.5 µm diameter and <10 nm height, consistent with previous reports of cellulose II nanocrystals [40]. A substantial fraction of cellulose II may prevent the crystals from adopting the typical needle shape commonly observed for cellulose Iβ nanocrystals [41]. We explore the implications of this mixed polymorph composition for CBM binding below.

The CNFs, prepared by mechanical disintegration of softwood pulp fiber, had a CI of 46%. This is on the low end of the typical CI of 45–65% reported for other solely mechanically treated CNFs [42, 43]. However, these previous reports of CNF crystallinity index were calculated using the Segal method (see ***Figure S6***), which often overestimates cellulose crystallinity [34]. AFM imaging of the CNFs showed a substantial amount of µm-scale fibers with diameters of 0.5–2 µm and tenths of µm lengths, as well as smaller fibrils with diameters on the 50–100 nm scale (***Figure S7***).

### *Sc*CBM2 and *Ba*CBM2 display distinct binding modes on insoluble polymers

We evaluated the binding efficiency of *Sc*CBM2 and *Ba*CBM2 on PET, chitin, and cellulose using pull-down assays. A Langmuir model was used to estimate binding affinity, *K*_d_, and binding capacity *B*_max_ (***Table 2***). We refer to binding efficiency as the relationship between binding affinity and capacity, which can be expressed by the partitioning coefficient *K*_r_ = *B*_max_∕*K*_d_, where both increased affinity (lower *K*_d_) and increased binding capacity lead to increased binding efficiency [9]. *K*_r_ can compare binding efficiency between different proteins on the same substrate, or between different substrates for a given protein. We stress that although the units of *K*_r_ are essentially meaningless, the magnitude of *K*_r_ depends on the units used to express *B*_max_, meaning the values are not always comparable.

**Table 2.**
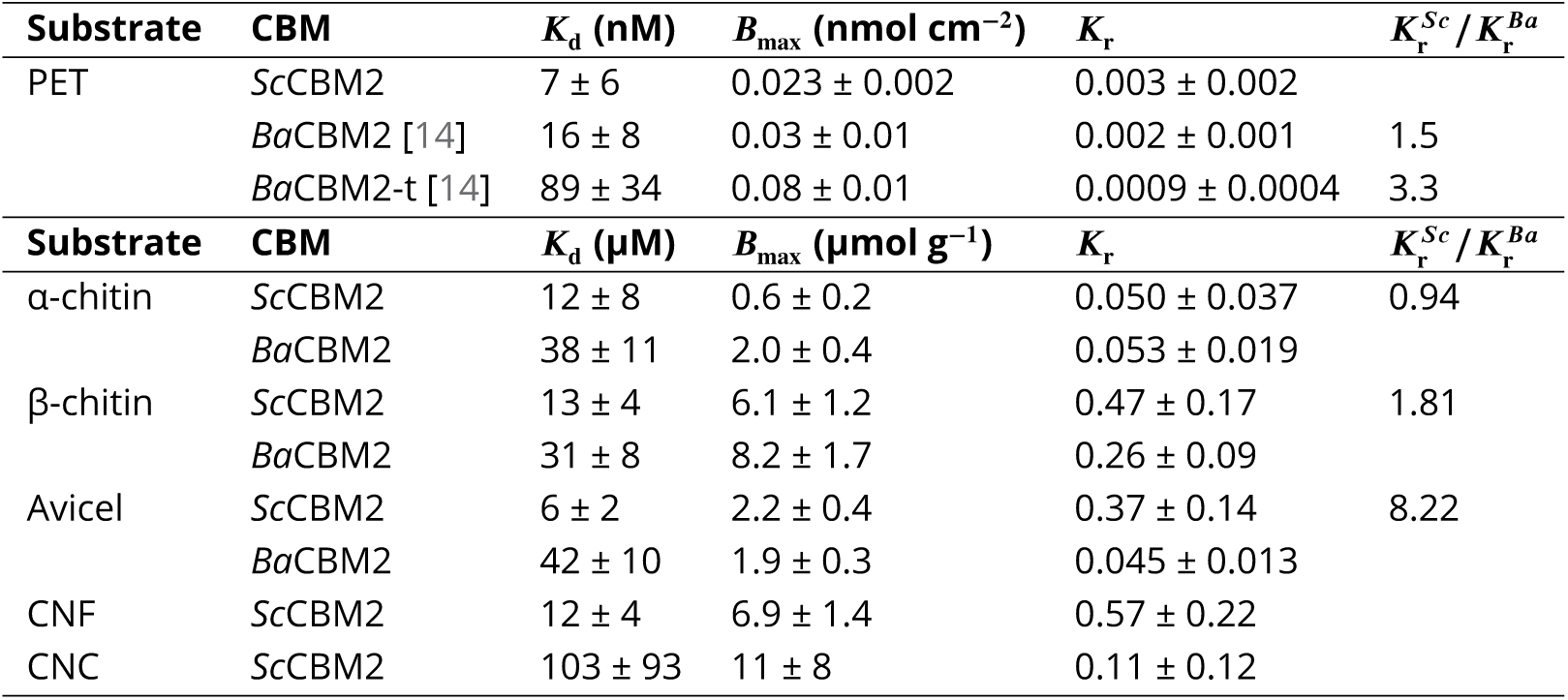
Dissociation constants (*K*_d_), binding capacities (*B*_max_), and partitioning coefficients (*K*_r_) of *Sc*CBM2 and *Ba*CBM2 on specified substrates. The ratio of partitioning coefficients [inlin] for *Sc*CBM2 and *Ba*CBM2 is shown for substrates where binding data are known for both proteins. We obtained binding data for *Ba*CBM2 and *Ba*CBM2-t on PET from Rennison *et al.* [14].

| Substrate | CBM | $K_d$ (nM) | $B_{\max}$ (nmol cm<sup>-2</sup>) | $K_r$ | $K_r^{Sc}/K_r^{Ba}$ |
| --- | --- | --- | --- | --- | --- |
| PET | ScCBM2 | 7 ± 6 | 0.023 ± 0.002 | 0.003 ± 0.002 |  |
|  | BaCBM2 [14] | 16 ± 8 | 0.03 ± 0.01 | 0.002 ± 0.001 | 1.5 |
|  | BaCBM2-t [14] | 89 ± 34 | 0.08 ± 0.01 | 0.0009 ± 0.0004 | 3.3 |
| Substrate | CBM | $K_d$ (μM) | $B_{\max}$ (μmol g<sup>-1</sup>) | $K_r$ | $K_r^{Sc}/K_r^{Ba}$ |
| α-chitin | ScCBM2 | 12 ± 8 | 0.6 ± 0.2 | 0.050 ± 0.037 | 0.94 |
|  | BaCBM2 | 38 ± 11 | 2.0 ± 0.4 | 0.053 ± 0.019 |  |
| β-chitin | ScCBM2 | 13 ± 4 | 6.1 ± 1.2 | 0.47 ± 0.17 | 1.81 |
|  | BaCBM2 | 31 ± 8 | 8.2 ± 1.7 | 0.26 ± 0.09 |  |
| Avicel | ScCBM2 | 6 ± 2 | 2.2 ± 0.4 | 0.37 ± 0.14 | 8.22 |
|  | BaCBM2 | 42 ± 10 | 1.9 ± 0.3 | 0.045 ± 0.013 |  |
| CNF | ScCBM2 | 12 ± 4 | 6.9 ± 1.4 | 0.57 ± 0.22 |  |
| CNC | ScCBM2 | 103 ± 93 | 11 ± 8 | 0.11 ± 0.12 |  |

*Ba*CBM2 binds PET with high affinity (***Figure 2***), and its affinity is strongly temperature-dependent [14]. We therefore measured PET binding of *Sc*CBM2 at 60°C, a more operationally relevant temperature that approaches the glass transition temperature, *T*_g_, of PET [44]. To enable detection at low concentrations, we expressed the protein with a GFP label. Interestingly, *Sc*CBM2 binding to crystalline PET was highly dependent on buffer composition (***Figure 2***). In Rennison *et al.*, binding of *Ba*CBM2 and *Ba*CBM2-t to PET was measured in 10 mM HEPES, 150 mM NaCl, 10 % glycerol at pH 7.5 (HEPES buffer). Using the same protocol for *Sc*CBM2, we observed very weak binding, and we could not fit the data to obtain binding parameters. This shows that *Sc*CBM2 exhibits negligible PET binding under these conditions.

**Figure 2.**
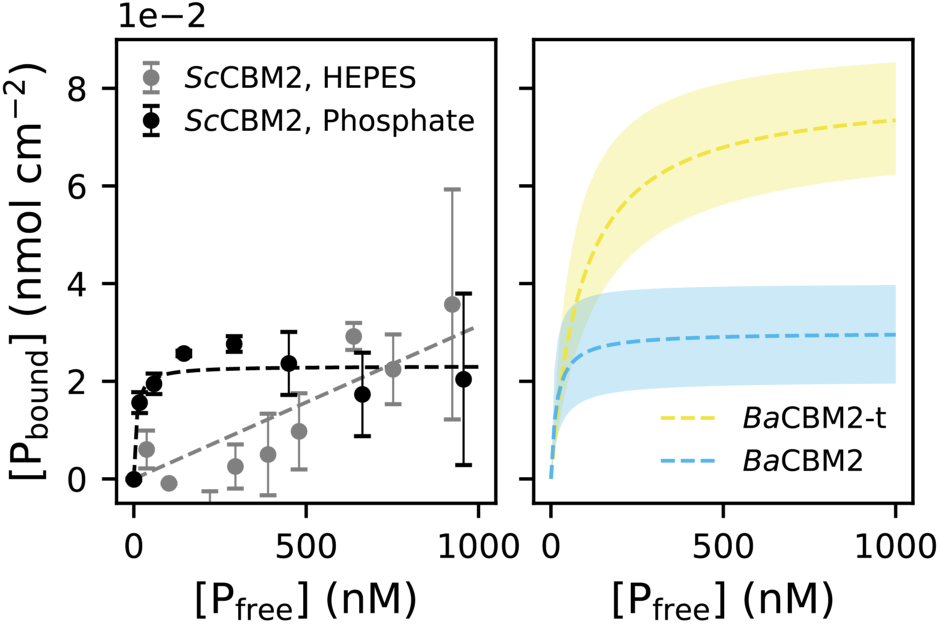
Binding of GFP-*Sc*CBM2 (left) on crystalline PET. Fitting to a Langmuir model (dashed lines) gave *K*_d_ = 7 ± 6 nM and *B*_max_ = 0.023 ± 0.002 nmol cm^−2^ in 50 mM sodium phosphate (pH 7.5) (black dots). Binding parameters could not be estimated in 10 mM HEPES, 150 mM NaCl, 10 % glycerol (pH 7.5) (grey dots). For comparison, binding data for *Ba*CBM2 (right) is shown, with *K*_d_ = 16 ± 8 nM and *B*_max_ = 0.03 ± 0.01 nmol cm^−2^ at 60°C, and the C-terminal truncated version *Ba*CBM2-t, *K*_d_ = 89 ± 34 nM and *B*_max_ = 0.08 ± 0.01 nmol cm^−2^ at 70°C, from Rennison *et al.* [14].

Glycerol is often added to stabilize proteins in solution, but in some cases it may inhibit protein-substrate interactions through direct competitive binding at the binding site and by modifying the protein’s hydration shell [45, 46]. High salt concentrations may also affect CBM binding by weakening electrostatic interactions [47]. Additionally, we speculated that SDS washing of the PET substrate before the assay may have led to adsorption of the anionic surfactant onto the PET film [48], which could negatively affect binding to PET [49, 50]. We therefore repeated the binding experiment using a simpler 50 mM sodium phosphate buffer at pH 7.5, and washed the PET film with ethanol solution instead of SDS. In this assay, *Sc*CBM2 displayed strong PET binding, with an estimated *K*_d_ of 7 ± 6 nM (***Figure 2***), similar to the previously reported value for *Ba*CBM2-t [14]. Increased binding affinity in a glycerol-free, low-ionic-strength buffer suggests a PET-binding mechanism that depends strongly on surface hydration and charge, whereas *Ba*CBM2 may be less affected by these factors. The observed effect is intriguing, but because several experimental variables were changed simultaneously, the data do not allow identification of the underlying cause.

The binding capacity of *Sc*CBM2 on PET is four times lower than that of the truncated *Ba*CBM2-t on the same crystalline PET film [14] (***Figure 2***) (***Table 2***). This indicates that *Ba*CBM2-t can access additional PET sites that are unavailable to both *Sc*CBM2 and full-length *Ba*CBM2. To test if this difference could be explained by size differences, we estimated the areas of the flat binding surfaces and the total footprint of the full-length and truncated *Ba*CBM2 versions using their AlphaFold structures (see ***Figure S8***). With estimated binding surface areas of 6.5 and 6.4 nm^2^ and total footprints occupying 8.7 and 8.3 nm^2^ for *Ba*CBM2 and *Ba*CBM2-t, respectively, the size difference is not sufficient to explain the large difference in binding capacity. Weber *et al.* showed by MD simulations that PET binding by *Ba*CBM2-t occurs in two steps [13]: An initial π-stacking of W9 and W44 with the phenyl rings of PET was followed by a conformational change that enables binding of W63 and N64. The missing β-strand in *Ba*CBM2-t (***Figure S4***-B) may confer increased flexibility, enabling this conformational change to adapt the binding site to heterogeneous PET structures.

For the natural substrates, chitin (*Ba*CBM2) and cellulose (*Sc*CBM2), we performed binding assays at room temperature to mimic their biological environments (***Figure 3***). Although these binding curves did not reach full saturation, they are sufficient for relative comparison of the two CBMs. Both CBMs bound to α-chitin and β-chitin with micromolar affinity, with *Ba*CBM2 exhibiting a 2-3-fold higher *K*_d_ for both chitin substrates. For α-chitin, the lower affinity of *Ba*CBM2 is compensated by a threefold increase in binding capacity, yielding similar overall binding efficiency (***Table 2***). This shows that *Ba*CBM2 accesses additional binding sites on α-chitin that are not accessible to *Sc*CBM2. Both CBMs showed much higher binding efficiency on β-chitin. The binding capacity on α-chitin was 10-fold lower for *Sc*CBM2, and 4-fold lower for *Ba*CBM2. To relate binding capacity to available surface area, we measured the particle size distributions of both chitin substrates by laser diffraction. Interestingly, they had very similar particle sizes with a volume-weighted mean diameter of 243 ± 4 µm (α-chitin), and 246 ± 5 µm (β-chitin), corresponding to a specific surface area of about 48 m^2^/kg for both substrates. Thus, the higher accessibility on β-chitin does not come from a higher surface area, but rather from structural or chemical differences on the substrate surfaces. Preference for either chitin polymorph has been shown for several chitin-binding CBMs and has been attributed to specific hydrogen bond interactions [51, 52].

**Figure 3.**
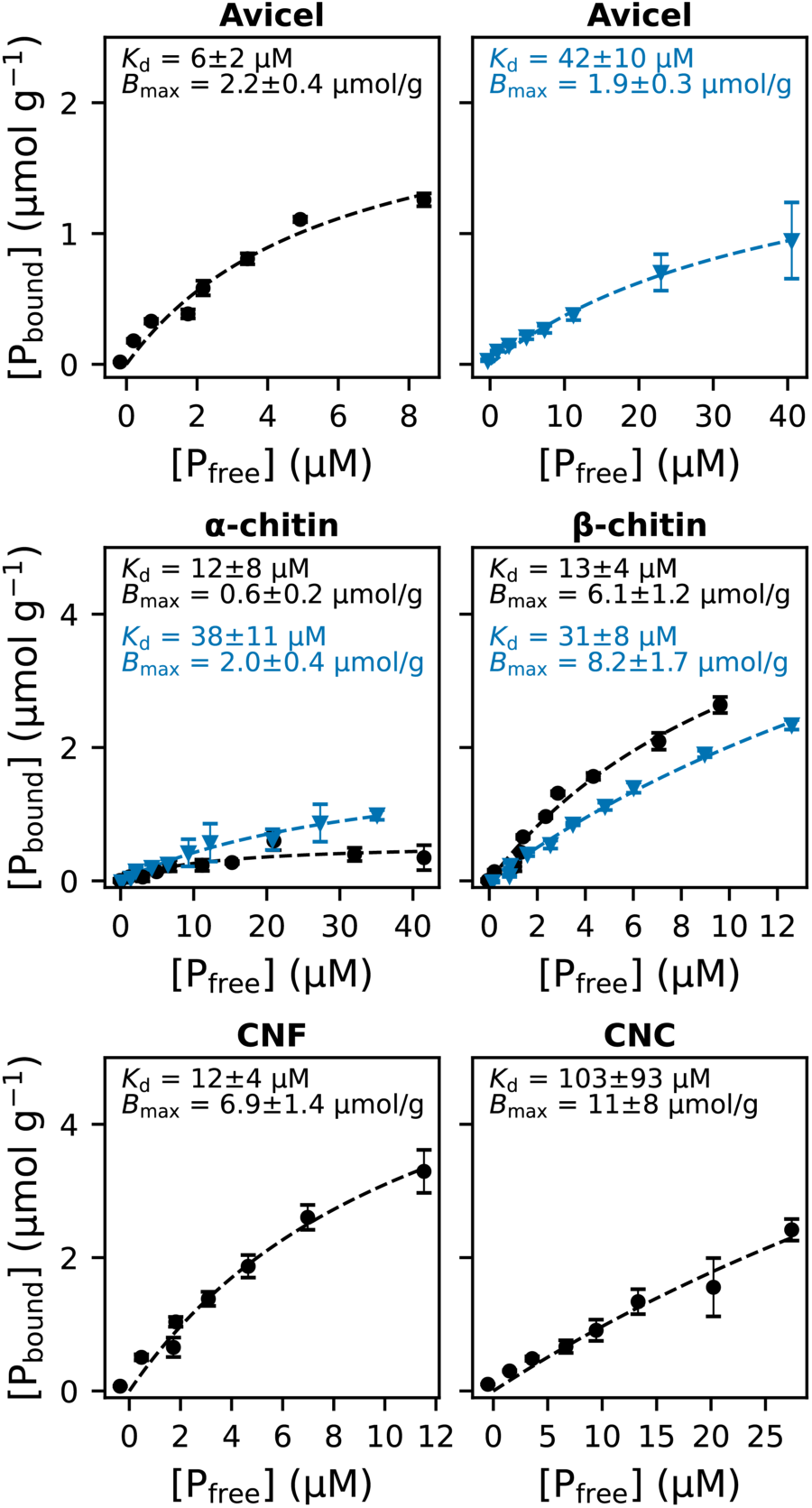
Binding of *Sc*CBM2 (black circles) and *Ba*CBM2 (blue triangles) to microcrystalline cellulose (Avicel), α-chitin, β-chitin, cellulose nanofibrils (CNF), and cellulose nanocrystals (CNC). The data were fit to a Langmuir model (dashed lines), and the estimated *K*_d_ and *B*_max_ are indicated on the plots.

As expected for its natural substrate, the affinity of *Sc*CBM2 to cellulose (6 ± 2 µM) was consistent with the previously published binding affinity of *Sc*LPMO10C (7.8 ± 1.0 µM) [53] and an order of magnitude higher than that of *Ba*CBM2, whereas the binding capacity was around 2 µmol g^−1^ for both proteins. Thus, in contrast to what was observed for chitin and PET, the eightfold higher *K*_r_ of *Sc*CBM2 on Avicel (***Table 2***) is a result of stronger binding rather than increased substrate accessibility.

The pull-down assays highlight key distinctions between the two CBMs. They are both efficient chitin-binders, but chitin is more accessible to *Ba*CBM2. Their differential binding to the α and β polymorphs of chitin shows that both prefer binding to β-chitin by accessing a higher number of binding sites. Next, *Sc*CBM2 is a much stronger cellulose-binder. Additionally, they display nanomolar affinity to PET, though *Sc*CBM2’s PET-binding ability depends strongly on the buffer environment. Interestingly, they show similar surface coverage on cellulose, PET, and β-chitin, indicating that they target equivalent binding sites with different affinities.

### *Sc*CBM2 binding depends on cellulose morphology

As the PET-binding ability of *Ba*CBM2 has been extensively characterized previously [14], we focused on investigating *Sc*CBM2 binding to a broader range of cellulose substrates. We include binding isotherms from pull-down assays with CNF and CNC in ***Figure 3***. The *B*_max_ values increase with the specific surface area of the cellulose particles; microcrystalline Avicel showed the lowest value, and CNC exhibited fivefold higher capacity per gram of substrate.

Comparison of the *K*_d_ values for Avicel and CNF revealed a clear correlation between crystallinity and affinity. *Sc*CBM2 binds to Avicel (CI = 85%) with twofold higher affinity than CNF (CI = 46%). This indicates that *Sc*CBM2 prefers to bind crystalline regions of cellulose and has a lower affinity for amorphous regions, which are more prevalent in the nanofibrils. This is consistent with the generally accepted binding mode of type A CBMs [1], and aligns with the reported preference of *Ba*CBM2 for high-crystallinity PET [14]. Still, the binding efficiency was higher for CNF than for Avicel 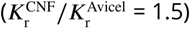 due to the greater binding capacity of the nanofibrils.

The CNC substrate could not be saturated under the conditions used in the pull-down assay (***Figure 3***); therefore, the *K*_d_ and *B*_max_ presented for CNC should be regarded as approximate. The affinity of *Sc*CBM2 is markedly lower for this substrate than for Avicel and CNF, despite its high crystallinity (91%). This reduced affinity resulted in substantially reduced binding efficiency on CNC 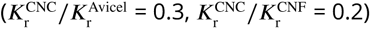. We propose that this reduction may be attributed, at least in part, to a suboptimal nanocrystal morphology, considering their relatively high cellulose II content (35%) (***Table 1***).

The crystallinity index obtained from XRD represents the bulk material and does not capture surface-specific features. The poor binding of *Sc*CBM2 to CNC (***Table 2***) suggests that cellulose II may be enriched at the crystal surface, where CBM binding occurs. Conversion to cellulose II disrupts the native cellulose Iβ structure, producing not only a distinct crystal polymorph but also disordered amorphous regions at the interfaces between cellulose II and cellulose Iβ, and both features could hinder CBM binding. In a recent study, Yang *et al.* used AFM single-molecule force spectroscopy to measure the adhesion forces of a family 3 CBM on cellulose II and cellulose Iβ [10]. They observed stronger binding to cellulose Iβ nanocrystals than to cellulose II and proposed that disrupting hydrogen bonds in cellulose II leads to misalignment of the hydrophobic face of the lattice (see ***Figure S10***) [37]. This disruption may prevent aromatic residues on the planar CBM surface from forming favorable interactions with the hydrophobic face of cellulose [7, 54, 55]. For this reason, *Sc*CBM2 may preferentially bind the flatter hydrophobic surface of cellulose Iβ, which could explain its reduced affinity to our CNC substrate. Its high dependence on cellulose morphology makes *Sc*CBM2 a potential candidate for cellulose characterization using CBMs [56].

### MD simulations reveal differences in binding surface water density

Distinct binding specificity patterns among type A CBMs have been attributed to differences in surface properties, including electrostatic potential and hydrophobicity [21]. Because these surface properties also modulate water organization at the protein-substrate interface, differences in surface hydration may contribute to the observed binding preferences. For example, Sahihi *et al.* observed a 24% reduction in enzyme-water hydrogen bonds upon adsorption of a cutinase onto PET, demonstrating that PET binding entails substantial dehydration of the binding interface [57]. Moreover, Yang *et al.* found that water molecules mediate surface interactions between CBM3 and cellulose, either by competing for binding sites or by reducing van der Waals forces [10].

To probe the effect of water density on binding, we carried out several 250-ns simulations of *Sc*CBM2 and *Ba*CBM2 in water, and observed that the water density on the surface of *Ba*CBM2 is consistently lower than that of *Sc*CBM2 (***Figure 4***). The radial distribution functions (***Figure 4***-A) show that the surface of *Sc*CBM2 is more hydrated than *Ba*CBM2. For clarity, the noise at *r <* 0.3 nm is due to the protein’s excluded volume, and the region 0.3 *< r <* 2.5 describes surface hydration. After *r* = 2.5, both RDFs converge to 1, which represents the bulk water, where the proteins do not affect water density. Water density maps on the protein surfaces (***Figure 4***-B) show an even distribution of water on the binding surface of *Sc*CBM2, whereas *Ba*CBM2 exhibits lower hydration around the first two aromatic residues, W9 and W44. If binding is mediated by hydration layers, lower water density implies that a smaller entropic dehydration reward is available for *Ba*CBM2 upon binding of these aromatic residues. Following this idea, protein surface hydration could affect binding affinity by modulating both this dehydration reward and the strength of hydrogen bonding and van der Waals interactions. We hypothesized that amino acid substitutions on the binding surface could modify the surface water density of *Sc*CBM2. In addition to the obvious W→H substitution on the surface of *Sc*CBM2 vs *Ba*CBM2, we noticed distinctly high water density around E10 of *Sc*CBM2 compared to T6 of *Ba*CBM2 (see ***Figure S11***), which prompted further investigation of the *Sc*CBM2-E10T mutant. Simulations of the E10T, H69W, and E10T-H69W variants in water (***Figure S12***) resulted in water RDF profiles that more closely resembled those of *Ba*CBM2. Specifically, both mutations appeared to reduce the RDF values in the range 0.3 *< r <* 1.5 relative to the wild type. These results demonstrate an interesting interplay between amino acid composition and surface hydration: while specific sequences or amino acid chemistries can not rationalize CBM2 specificity, they may indirectly affect selectivity and binding strength by tuning surface hydration.

**Figure 4.**
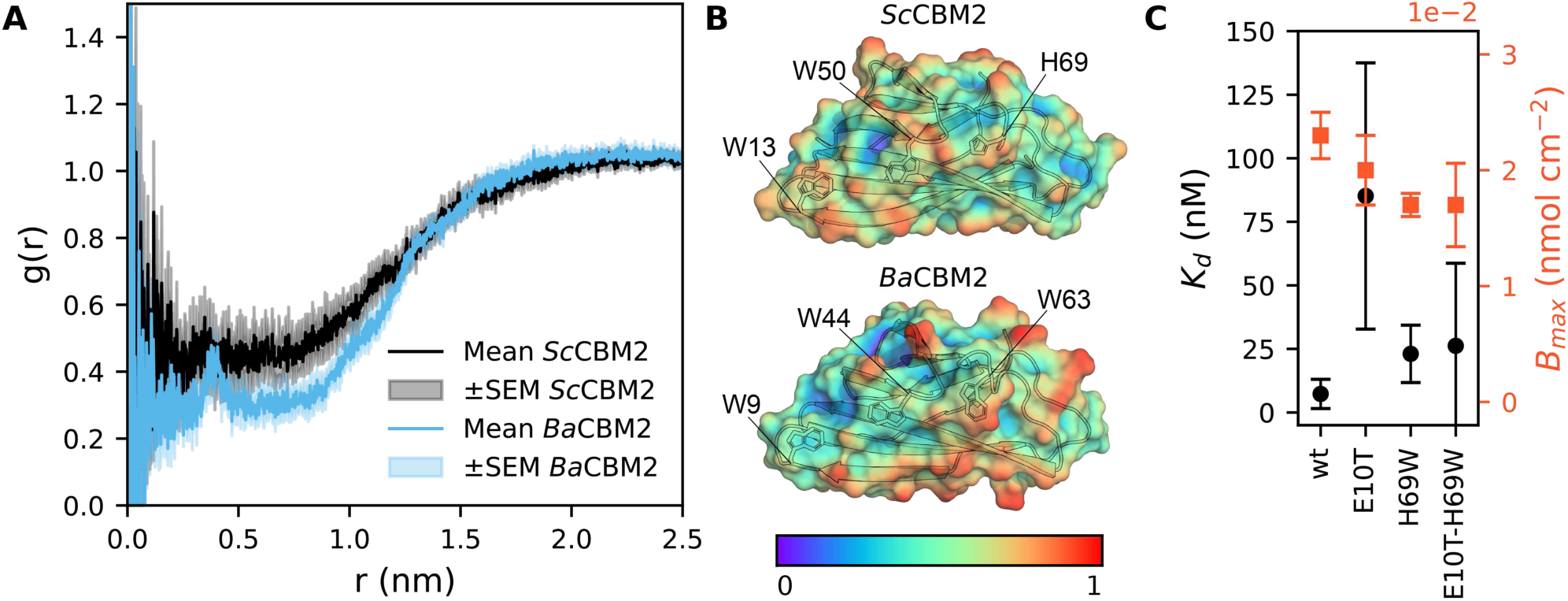
(A) Radial distribution functions (RDF) showing the density of water molecules *g*(*r*) as a function of distance *r* from the protein’s center of mass during the 250 ns simulation of *Sc*CBM2 and *Ba*CBM2 in water. The error bars show the standard error of the mean (SEM) between the RDFs calculated from 5 × 50 ns blocks during the 250 ns simulations. (B) Surface water density on *Sc*CBM2 (upper) and *Ba*CBM2 (lower), where 0 and 1 represent minimal and maximal water density, respectively. (C) Experimental *K*_d_ and *B*_max_ values for *Sc*CBM2 (wt and variants) on PET. Full binding curves are shown in ***Figure S13***.

Based on the simulations, we produced GFP-labeled E10T, H69W, and E10T-H69W variants of *Sc*CBM2 to assess the effect of these substitutions on PET binding. All three variants displayed slightly poorer PET binding than wild-type *Sc*CBM2 (***Figure 4***-C and ***Figure S13***). The H69W substitution caused a modest reduction in binding capacity and a threefold increase in *K*_d_. The E10T mutant exhibited the lowest apparent affinity, with a *K*_d_ of 85 ± 52 nM. This result is consistent with the possibility that reduced water density around W13 may weaken its interactions with PET. Interestingly, the E10T-H69W double mutant displayed binding parameters very similar to those of the H69W variant, suggesting that the contribution of the H69W substitution dominates in tuning *Sc*CBM2 binding efficiency. Together with the limited sensitivity of the fluorescence assay at low protein concentrations, the dominating effect of the H69W substitution prevents us from elucidating how the E10T substitution affects PET binding. Still, these results suggest that modifying surface hydration through simple amino acid substitutions can tune *Sc*CBM2 adsorption on PET. If so, reduced surface hydration negatively impacts affinity, which aligns with a binding mechanism driven by favorable removal of water from the protein-substrate interface.

### Surface solvent accessibility assessed by paramagnetic relaxation enhancement

After observing that sequence and fold alone cannot explain CBM2 substrate specificity, we decided to explore less obvious properties. CBM2 modules are often considered as rigid proteins. However, the PET-binding mode simulated by Weber *et al.* shows conformational flexibility of *Ba*CBM2-t [13], which inspired us to assess protein dynamics in relation to substrate binding using NMR. NMR is a powerful tool for exploring the dynamic properties of proteins in solution, and nuclear spin relaxation rates provide information about local flexibility and exchange dynamics [58, 59].

We measured backbone amide ^15^N and ^1^H longitudinal (*R*_1_) and transverse (*R*_2_) relaxation rates for *Sc*CBM2 and *Ba*CBM2 (***Figure S14***). The relatively small variations in *R*_1_ and *R*_2_ are consistent with the overall rigidity of both proteins. However, the aromatic triad of *Sc*CBM2 exhibited higher relaxation rates than the rest of the protein, suggesting that this region has reduced backbone flexibility on the picosecond-to-nanosecond timescale. By contrast, the corresponding residues in the aromatic triad of *Ba*CBM2 exhibited lower relaxation rates, consistent with higher flexibility. Our MD simulations showed differences in water distribution on the surfaces of *Sc*CBM2 and *Ba*CBM2, and *Sc*CBM2 variants with lower surface hydration showed weaker binding to crystalline PET. To compare the solvent accessibility of *Ba*CBM2 and *Sc*CBM2, we measured solvent paramagnetic relaxation enhancement (sPRE) using the Gd^3+^ agent gadodiamide (***Figure S15***). In the presence of this paramagnetic probe, ^1^H relaxation rates increase linearly with increasing gadodiamide concentration, and the sPRE is obtained as the slope of this relationship [60, 61]. The magnitude of the relaxation enhancement depends on the proton’s distance from the probe. Thus, sPRE can visualize solvent accessibility, provided the probe does not directly bind to the protein.

Both CBMs exhibited high sPRE around the aromatic triad, but the values were significantly higher near W44/W63 in *Ba*CBM2 than near W50/H69 in *Sc*CBM2 (***Figure 5***-A). Interestingly, this sPRE trend does not match the simulated water densities (***Figure 5***-B). In *Ba*CBM2, the combination of high sPRE and low simulated water density suggests that fewer immobilized water molecules shield the aromatic triad from the bulk solvent. Conversely, *Sc*CBM2 displayed lower sPRE despite higher simulated water density. This suggests that the aromatic residues are less accessible from the bulk solvent because of a denser hydration layer on the binding surface. Taken together, these results point to a larger contribution from entropically favorable dehydration available for *Sc*CBM2 upon binding of the aromatic triad. The observed denser hydration layer is consistent with our hypothesis on how surface water density may affect affinity, because the more hydrated *Sc*CBM2 consistently shows higher affinity for cellulose, chitin, and PET (***Table 2***).

**Figure 5.**
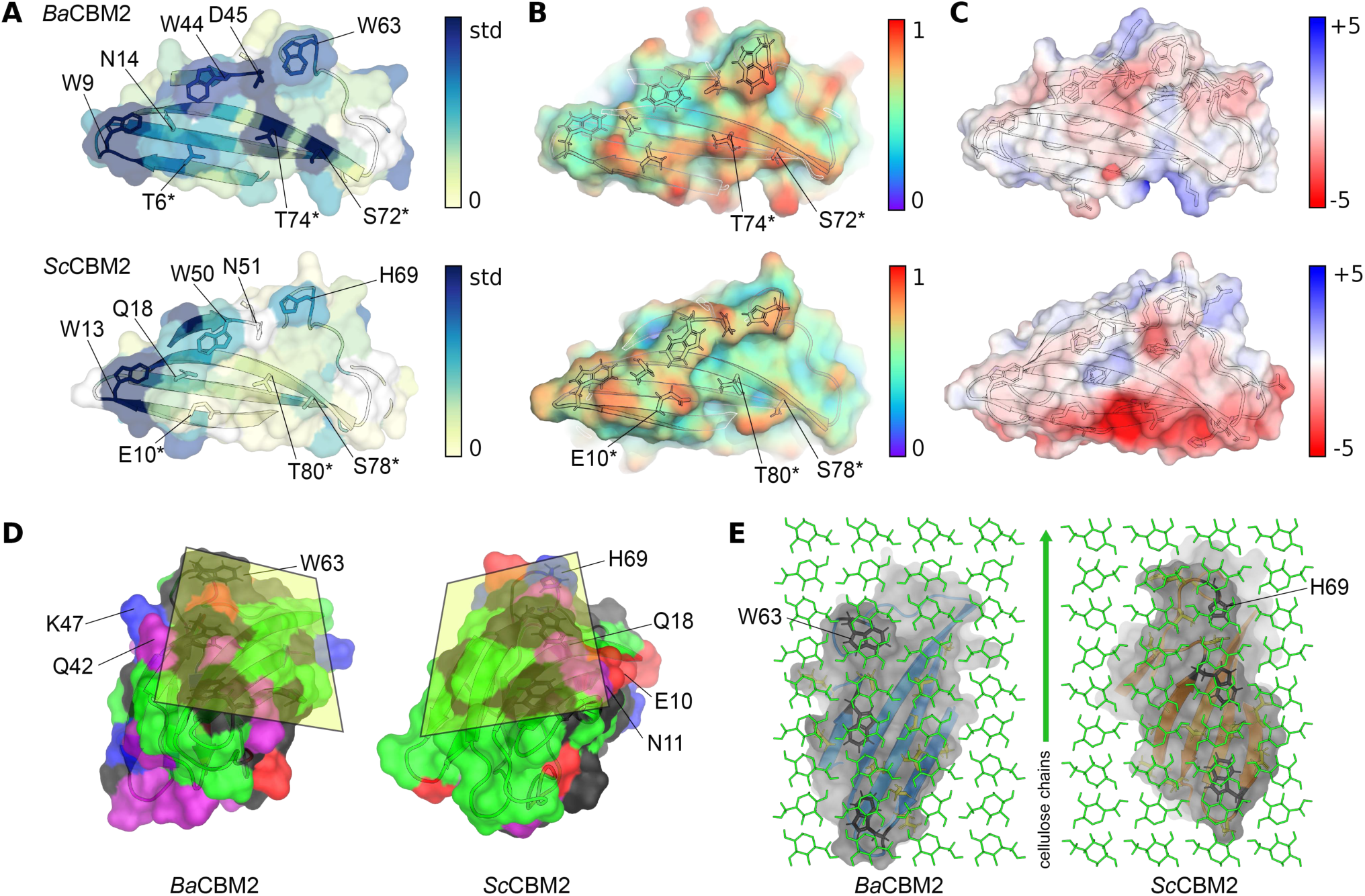
Surface properties mapped on *Ba*CBM2 (upper panel) and *Sc*CBM2 (lower panel) structures. (A) Experimental ^1^H *R*_2_ sPRE measured for backbone resonances on AlphaFold [31] structures (see *Figure S15*). The coloring represents the relative sPRE from 0 (light yellow) to one standard deviation above the mean sPRE (std, dark blue). Residues on the main β-sheet surface with significantly different sPRE between the two CBMs are shown as sticks and labeled with an asterisk (*). Residues with high sPRE within and between aromatic triads are shown as sticks. ^1^H *R*_2_ sPRE for N51 of *Sc*CBM2 could not be determined, but high ^1^H *R*_1_ sPRE was measured for this residue (*Figure S15*), indicating a similar solvent accessibility as for the corresponding D45 of *Ba*CBM2. (B) Water density around the aromatic triad from the MD simulations. (C) Electrostatic potential surfaces (created using the APBS plugin in PyMOL [62]) on AlphaFold [31] structures, with charged and aromatic residues shown as sticks. (D) Simulated structures showing the proposed binding interface planes on the right- (*Ba*CBM2) and left-hand (*Sc*CBM2) side of the aromatic triad. Residues are colored by their chemical properties: polar = green, neutral = purple, basic = blue, acidic = red, and hydrophobic = black. (E) Illustration of proposed binding poses, showing interaction of the aromatic triad (black sticks) with a single cellulose chain and polar residues (yellow sticks) available for hydrogen bonding with adjacent chains, at the hydrophobic face of a cellulose crystal (green sticks). The surface representations show similar footprints of the suggested binding sites.

On the main β-sheet surface containing the first two tryptophans of the aromatic triad, the relative solvent accessibility (sPRE) around S72 and T74 of *Ba*CBM2 was higher than around the corresponding residues, S78 and T80, of *Sc*CBM2. This result is consistent with our simulations, which show higher water density in this region for *Ba*CBM2 (***Figure 5***-B). From the simulated solution structures, the hydrophobic strip of *Sc*CBM2 protrudes up and away from the lower end of the β-sheet, placing S78 and T80 in a groove almost perpendicular to the binding face (***Figure 5***-B). This groove is formed by the bulky residues E10, N11, and Q18, which could explain why the polar residues on the lower end of the β-sheet are less solvent-exposed. In *Ba*CBM2, the corresponding residues, S72 and T74, are more aligned with the aromatic triad and may therefore contribute more directly to substrate binding. We discuss the impact of these structural differences on CBM binding orientation later.

### Comparison of surface electrostatics

We next assessed differences in surface electrostatic potential between the CBMs. At neutral pH, the net charges of *Ba*CBM2 and *Sc*CBM2 are -1.5 and -6.7, respectively. Both CBMs display a mostly neutral electrostatic potential along the aromatic triad (***Figure 5***-C). *Sc*CBM2 is more negatively charged overall and carries several negatively charged residues on the main β-sheet, whereas *Ba*CBM2 has polar uncharged residues in these positions, giving it a more neutral electrostatic potential on the β-sheet surface.

Electrostatic potential has been shown to affect PET hydrolase activity. PET hydrolases typically target PET through non-specific interactions involving aromatic, aliphatic, and polar uncharged residues [57, 63, 64]. Although PET binding by these enzymes is dominated by hydrophobic interactions, π-stacking, and hydrogen bonding, surface charges far from the active site strongly influence activity. Introducing positive charges can enhance PET binding through electrostatic attraction to amorphous PET, while adding negative patches away from the active site can increase enzymatic activity by directing binding towards a catalytically productive orientation [64–66]. Similarly, the negative charge across the β-sheet of *Sc*CBM2 may play a role in directing its orientation on polymer substrates through electrostatic repulsion.

The binding assays (***Figure 2***) showed that *Sc*CBM2 binding to PET is highly sensitive to buffer composition, indicating that surface hydration and charged residues strongly affect *Sc*CBM2 binding to PET. In addition to weakening electrostatic repulsion and hydrogen-bond interactions, high salt concentrations modify the arrangement of water molecules in the hydration layer covering polymer surfaces [67]. As discussed above, *Sc*CBM2 exhibits higher structural water density at the aromatic triad than *Ba*CBM2, implying a larger entropic gain upon displacing immobilized water during binding to W13 and W50. Charged residues surrounding hydrophobic binding sites often aid in clearing water from the binding site [22], and this effect may be weakened at higher ionic strength. We hypothesize that *Sc*CBM2’s dependence on buffer composition arises from the combined effects of charged residues in both displacing water and directing binding toward the hydrophobic aromatic triad. However, we cannot exclude that anionic surfactant residues from the SDS washing step of the PET film may have increased the negative charge on the PET surface, potentially leading to excessive electrostatic repulsion for the more negatively charged *Sc*CBM2 than for *Ba*CBM2.

### CBM2 orientation on crystalline polymers

The spacing of the aromatic rings in the CBMs (10.7–12.6 Å, ***Figure S16***) matches the distance between every other glucose ring in a crystalline cellulose chain (10.4 Å) [37], and each aromatic ring in crystalline PET (10.8 Å) [68]. This complementary geometry suggests a binding pose in which the planar strip engages with a single polymer chain, as suggested by Tormo *et al.* [69]. CBMs that preferentially or specifically target crystalline substrates often require additional contacts across adjacent chains, for example through hydrogen bonding [10, 70]. Previous studies show that the interaction between *Sc*CBM2 and cellulose is focused on the aromatic strip, but other residues also participate in binding [53]. Interestingly, these additional residues are located on the left-hand side (in ***Figure 5***-D) of the aromatic strip, in contrast to the binding pose of *Ba*CBM2-t on PET, where additional residues are located on the right-hand side (in ***Figure 5***-D; ***Figure S4***) [13].

Combining these observations with the fact that both CBMs display similar binding capacities on both cellulose and PET, we propose that they may target equivalent binding sites on crystalline substrates from different orientations (***Figure 5***-E), resulting in similar surface coverage (***Figure S8***). This would explain how substrate specificity is not determined by sequence and fold alone, but rather likely depends on the position of polar amino acid contacts and structural compatibility along the protein-substrate interface. From our structural comparison (***Figure 5***-D; ***Figure S4***), we propose that structural compatibility is determined by surface topology, and that bulky residues (E10, N11, and Q18 in *Sc*CBM2 and Q42 and K47 in *Ba*CBM2) and polar amino acids on either side of the aromatic triad determine its binding orientation. Following this logic, conformational flexibility could also influence specificity, because more flexible CBMs may structurally adapt to varying substrate surface topologies.

We rationalize the idea that similar CBM2 folds may bind to crystalline substrates with opposite orientations by considering the nature of each energetic contribution to the binding interaction, and the locations in which each of them dominates. For crystalline substrates, the most important contribution is dehydration of the protein-substrate interface, which is driven by entropically favorable removal of water molecules from hydrophobic surfaces. This implies that interaction is favored by binding areas with high water density and large surface areas. The location of regions with high water density matches the proposed binding surfaces: *Ba*CBM2 showed higher water density and sPRE on the lower end of the main β-sheet, whereas for *Sc*CBM2 the more hydrated regions were on the left-hand side (in ***Figure 5***-D) of the aromatic triad. This indicates that surface hydration not only affects affinity, but may also favor certain binding orientations, as regions with high water density would produce a larger entropy gain upon binding. The next contribution is π-stacking interactions (CH-π in cellulose and chitin, and π-π in PET), which dominate between the targeted polymer chain and the aromatic triad residues. These interactions are known to be orientationally promiscuous [71, 72]. In that sense, the CBM’s direction along the targeted polymer chain is not significant for obtaining stable π-stacking interactions. A similar principle was demonstrated by Georgelis *et al.*, who solved the crystal structure of a type A CBM EXLX1 bound to cellohexaose, where two CBM proteins bound to opposite faces of a single oligomer in a sandwich-like arrangement [73]. Since hydrophobic interactions and π-stacking permit multiple orientations, specific hydrogen-bonding with adjacent polymer chains and topological complementarity are likely the main determinants of the CBM’s binding pose. Finally, electrostatic repulsion may tune the orientation of CBMs that contain patches of strong negative or positive charge, depending on substrate properties.

We observed the largest affinity difference between the two CBMs on cellulose, where *Sc*CBM2 showed much stronger binding. In contrast, the affinity difference for PET was very low. Relative affinity differences between *Sc*CBM2 and *Ba*CBM2 on each substrate may reflect the relative contributions from polar interactions versus dehydration of the protein-substrate interface, where cellulose-binding seems to be more strongly influenced by the latter. For chitin and PET, which contain more polar regions and offer greater hydrogen-bonding potential, favorable hydrogen-bond interactions and electrostatics may play a more dominant role in the binding mechanism.

### Probing *Ba*CBM2 interactions with PET by NMR

Fusion of CBMs to PET-hydrolyzing enzymes can be used to tune affinity, activity, and thermostability [15–20]. Harnessing the full potential of CBMs for enzyme engineering requires a more detailed understanding of the molecular mechanism of substrate binding. *Ba*CBM2 is among the most well-characterized PET-binding CBMs to date [13, 14, 74, 75], making it a useful model CBM to explore mechanistic details of PET binding. To obtain residue-specific insight into *Ba*CBM2 binding to PET, we measured chemical shift perturbations (CSP, Δ*δ*) and changes in *R*_2_ relaxation rates and sPRE in the presence of PET nanoparticles and the soluble monomer bis(2-hydroxyethyl) terephthalate (BHET). CSP arise from changes in the chemical environment experienced by observed nuclei upon ligand binding or conformational rearrangement. For residues undergoing chemical exchange in the fast-exchange regime, where the exchange rate, *k*_ex_ = *k*_on_ + *k*_off_, is faster than the difference in resonance frequency *ω* between the two states (i.e., *k*_ex_ *>>* Δ*ω*), the observed chemical shift is the population-weighted average of the shifts in the free and bound state [59]. Longer-lived interactions may additionally cause line broadening through an increase in *R*_2_ relaxation rates due to chemical exchange (*R*_ex_) [76].

In the presence of BHET, we observed large CSP for residues of the aromatic triad (W9, W44, and W63) and for residues surrounding W9 (N8, G10, and S11) (***Figure 6***-A). These perturbations are consistent with BHET binding at all aromatic residues in the aromatic triad and suggest that the interaction is stronger or most extensively populated at W9, where several neighboring residues were also perturbed. Indeed, *R*_2_ relaxation rates were also slightly increased for W9, indicating slower exchange dynamics at the first tryptophan of the aromatic triad compared to the others. We also observed a decrease in sPRE for all three tryptophans in the presence of BHET, with the largest decrease at W9. This decreased sPRE shows that the aromatic triad is shielded from bulk solvent as a consequence of BHET binding. Because these NMR observables represent averages over all states present in the sample, the pronounced changes at W9, including increased *R*_2_ and decreased sPRE, are consistent with a greater population of states in which BHET is bound to W9. Additionally, we observed decreased sPRE and increased *R*_2_ for several surface-exposed hydrogen-bond donors far from the aromatic triad, including T2, K20, K28, K31, and N88. These changes suggest transient electrostatic interactions between these residues and the ester or hydroxyl groups of BHET.

**Figure 6.**
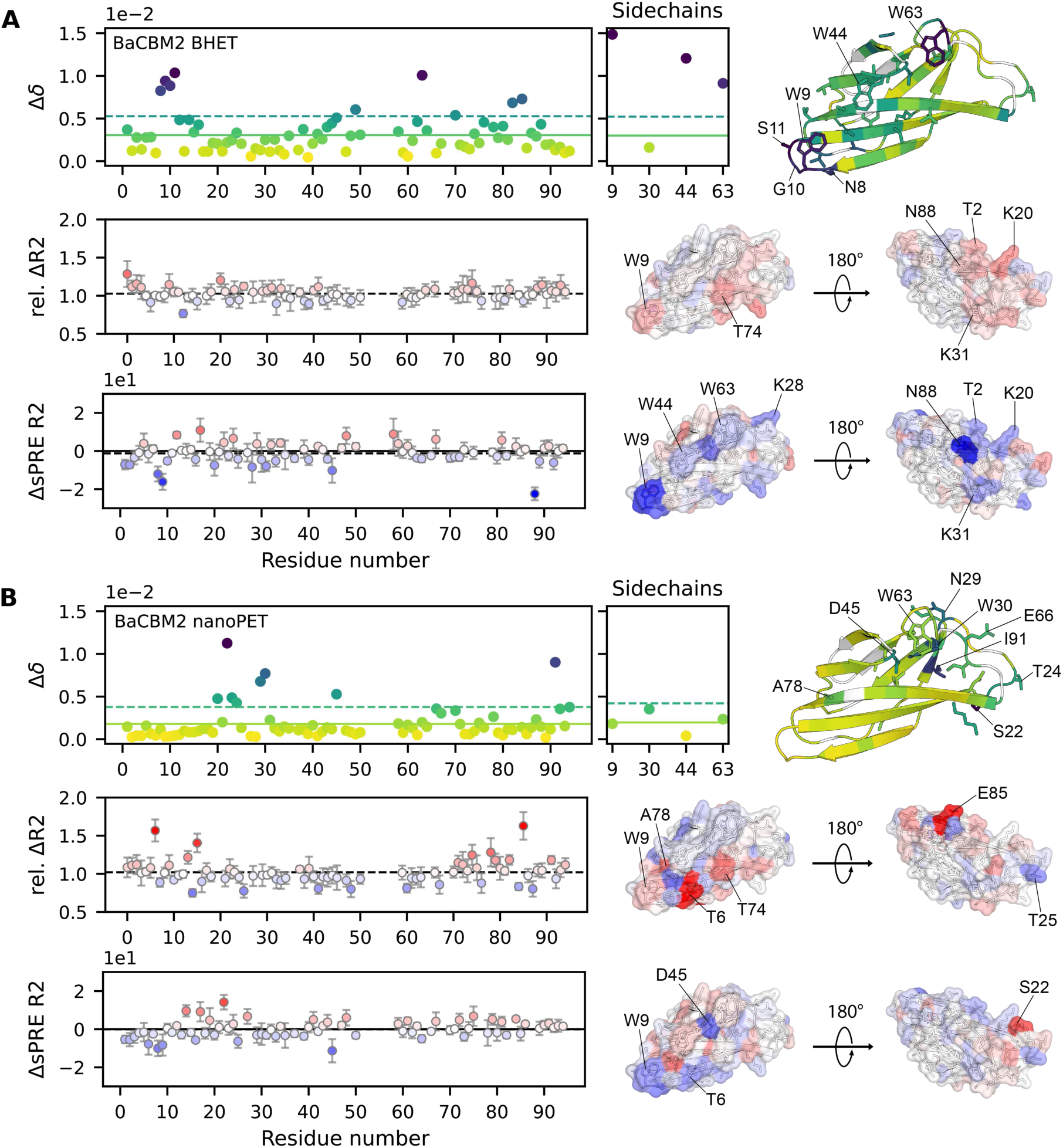
Chemical shift perturbations (Δ*δ*), relative change in *R*_2_ relaxation rate (Δ*R*_2_), and change in *R*_2_ sPRE (ΔsPRE) measured in ^15^N HSQC NMR spectra of *Ba*CBM2 in the presence of BHET (A) and PET nanoparticles (B). Data points are colored by magnitude and shown on the structures to the right. Residues with CSP above the mean are shown as sticks.

Using PET nanoparticles as substrate, CSP were not observed along the first two residues of the aromatic triad, W9 and W44 (***Figure 6***-B). Instead, CSP were most pronounced around W63 and the two loops that connect the β-sheets below W63 (e.g., around S22, T24, N29, W30, E66, and I91). In this region, we detected more positive changes in sPRE, indicating that these residues are more solvent-exposed in the ensemble of states that exist in the presence of PET. Additionally, the data indicate that a separate interaction process occurs around W9 and T6. Here, we observed a solvent-shielding effect, with lowered sPRE and increased *R*_2_ relaxation, but no significant CSP. These results indicate the presence of transient, low-population protein states involved in the PET binding mechanism of *Ba*CBM2.

Comparing the NMR data on soluble BHET with those on PET nanoparticles reveals differences in *Ba*CBM2’s dynamic interaction with each substrate. The absence of detectable CSP around W9 and W44 in the presence of PET does not mean that these residues are not involved in PET binding. NMR observables depend on the chemical environment of the observed nuclei, the exchange dynamics, and the populations of the states involved [77]. If binding produces a state with substantially higher *R*_2_ than the free protein, such as a stable nanoparticle-bound state, its signal may broaden beyond detectability. In the slow-exchange limit, where *k*_ex_ *<<* Δ*R*_2_, the bound-state magnetization decays before it can contribute to the observed signal, so the signal reflects only NMR-visible states. We observed a 6% decrease in the average signal intensity of *Ba*CBM2 in the presence of PET nanoparticles, which we did not observe in the BHET sample. The average *R*_2_ relaxation rate increased by 2.6 s^−1^ in the presence of PET. Because the increase in *R*_2_ was not uniform across all amino acids, it is unlikely to arise solely from a global lifetime line-broadening contribution, *R*_llb_, caused by formation of a high-molecular-weight complex that completely restricts rotational freedom of the CBM [78]. Instead, the reduced signal intensity indicates that a fraction of the protein is depleted from solution by binding to PET, where the fully bound state is NMR invisible. Thus, unlike BHET, the PET data cannot be interpreted using a simple two-state model consisting only of free and bound protein. The residue-specific changes in *R*_2_ may reflect chemical exchange (*R*_ex_) and partial restriction of motion (*R*_llb_) in transiently bound states [79].

In the BHET experiments, observed CSP largely correlated with increased *R*_2_ and decreased sPRE (***Figure 6***-A). This pattern is consistent with a system undergoing fast-intermediate exchange between two NMR-visible states, in which the interacting residues experience altered chemical environments, increased exchange broadening (*R*_ex_), and partial shielding from the bulk solvent.

Conversely, we measured minimal relaxation effects in the high-CSP regions of *Ba*CBM2 in the PET sample (***Figure 6***-B), and many of these residues (e.g., around S22) displayed increased sPRE, indicating higher solvent accessibility relative to the free protein. These data are consistent with a PET-induced conformational state that remains sufficiently populated and mobile to be observed by NMR, but they do not establish whether the state forms before binding, during binding, or after desorption. One possible interpretation is that, if *Ba*CBM2 binding involves a conformational change similar to that shown by MD simulations for *Ba*CBM2-t [13], the observed CSP could be related to a rearrangement back to the solution conformation after desorption. While the location of the high-CSP region around W63 aligns with this interpretation, it is not possible to confirm from our data whether such a conformational rearrangement takes place.

We propose that the increased *R*_2_ and decreased sPRE observed around W9 in the presence of PET (***Figure 6***-B) could arise from short-lived encounter complexes preceding productive binding. In such complexes, W9 and nearby residues, including T6, N8, and T74, may transiently approach the PET surface while sampling favorable orientations. Transient encounter states have been reported for proteins interacting with nanoparticles, and these states may involve interactions with the nanoparticle hydration layer rather than direct contact with the underlying surface [79–81]. If the encounter complex partially restricts the protein without fully immobilizing it, it could produce more local *R*_llb_ contributions to *R*_2_. Its low population and short lifetime could explain why it produces detectable relaxation and sPRE changes but no measurable CSP. The hydration layer on the PET nanoparticles is distinct from bulk solvent in a manner similar to the hydration layer on the protein itself. Transient encounters through this interface likely exclude bulk solvent, lowering the sPRE of residues at the encounter interface, consistent with our NMR data.

These observations do not uniquely distinguish specific protein states involved in the PET-binding mechanism of *Ba*CBM2. However, they highlight the added complexity of the dynamics involved in binding to PET nanoparticles compared to soluble BHET molecules. Whereas the BHET experiments show that specific interactions with phenyl rings are clearly concentrated on the aromatic triad, the PET experiments reveal binding regions other than the aromatic triad. This reflects the heterogeneity of the PET surface compared to the simple monomeric ligand BHET. Notably, the observed decrease in sPRE on the lower part of the β-sheet (residues 2-9) hints at their involvement in PET binding. This aligns with our previous suggestion that *Ba*CBM2-PET interactions are strengthened by polar contacts on the main β-sheet, and that the binding pose is similar to the simulated pose of *Ba*CBM2-t, though it may be less conformationally flexible giving it restricted access to structurally varied binding sites. Additionally, the CSP and sPRE data suggest that PET binding involves conformational or dynamic changes that are not observed in the presence of BHET. These insights challenge the commonly accepted view of CBM2 molecules as dynamically rigid proteins and call for more research on their ability to adapt to various substrates.

## Conclusion

CBM2 substrate specificity and selectivity clearly depend on a balance of multiple contributing factors. Despite analyzing hundreds of CBM2 sequences, we found that sequence conservation alone was insufficient to explain substrate specificity. We used two model CBM2s, *Sc*CBM2 and *Ba*CBM2, to investigate the impact of sequence, structure, surface hydration, and electrostatics on their binding modes to cellulose, chitin, and PET. *Sc*CBM2 showed higher surface water density around the aromatic triad and higher affinity to all of the tested substrates, suggesting that the location and density of hydrated patches on the protein surface may influence affinity for crystalline substrates. We demonstrated that surface hydration modulates CBM2 affinity to PET, supporting a mechanism driven by dehydration of the aromatic triad. Moreover, we showed how substrate crystallinity and morphology affect CBM binding. While the two CBMs have similar folds, we found significant differences in surface topology and position of polar and charged residues. Despite these differences, both CBMs showed the same surface coverage on cellulose and PET. This led us to propose that these CBMs may bind with opposite orientations to equivalent binding sites on crystalline substrates, where the aromatic triad aligns with a single polymer chain and polar contacts with adjacent chains determine their orientation. Our findings suggest that CBM2 substrate selectivity is governed by topological and hydrogen-bond complementarity between the protein’s planar binding site and the substrate’s surface, and not specifically by sequence and fold. The balance between primary hydrophobic and π-stacking interactions along the aromatic triad, secondary hydrogen bonding of polar residues with adjacent chains, and binding-site dehydration could be important for understanding CBM2 selectivity, promiscuity, and binding dynamics. Future studies of CBM2 binding modes could benefit from determining the relative weights of each of these contributions. This could potentially be used to classify both CBM proteins and substrates by dominant contributions, beyond classifications based on sequence, phylogeny, and fold.

## Experimental procedures

### Polymer substrates

Biaxially oriented PET film (0.1 mm thickness, 31.6% ± 2.2% crystallinity [14], ES30-FM-000200) was purchased from Goodfellow (UK). PET nanoparticles were produced as described previously [82]. In short, 300 mg of amorphous PET film (Goodfellow, UK) was dissolved in 20 mL hexafluoroisopropanol (HFIP) and incubated at rt overnight. The solution was then drip-fed into 300 mL of ultra-pure water using a burette at approximately 0.5 mL/min, under vigorous stirring. Large PET particles were removed by filtration, and the HFIP was removed under reduced pressure (200 mbar and 50°C). The nanoPET concentration was then determined by drying an aliquot at 105°C for 24 h, and particle size was measured using a Zetasizer Nano (Malvern, UK). Avicel PH-101 (11363) and α-chitin (from shrimp shells, C7170) were purchased from Sigma-Aldrich (Germany). β-chitin (< 500 µm particle size) was produced by mechanical grinding of β-chitin flakes purchased from Mahtani Chitosan (India) using a Culatti Micro-Fine Mill (IKA-Werke GmbH, Germany) with a 0.5 mm sieve. Cellulose nanofibrils (CNF) and cellulose nanocrystals (CNC) were kindly gifted from RISE PFI (Trond-heim, Norway). CNF were produced by mechanical pre-treatment of bleached softwood pulp and fibrillation by high-pressure homogenization as described previously [53]. CNC were produced by sulfuric acid hydrolysis of cellulose fibers as described in [83].

### X-ray diffraction (XRD)

XRD patterns were recorded on a D8 A25 DaVinci X-ray Diffractometer (Bruker AXS) with Cu Kα radiation, Bragg-Brentano geometry, with fixed 2.5° primary and secondary Soller slits. Data were collected from 10° to 75° 2θ, with a step size of 0.2° over 2 h. Rietveld fitting of XRD data was performed in TOPAS v6 (Bruker AXS). A 3rd order polynomial function was used to model the background. The crystalline fraction was modeled using the cellulose Iβ structure published by Nishiyama *et al.* [37], and, in the case of CNC, the cellulose II structure published by Kaduk in the ICCD database (PDF number 00-056-1717) [38]. Pseudo-Voigt line shapes were used in the fitting, and a (0 0 1) March-Dollase correction was applied for preferred orientation, as recommended by Ju *et al.* [39]. The amorphous fraction was modeled using Pseudo-Voigt line shapes fixed at the amorphous peak positions (20.52 and 38.87°) defined by Ju *et al.* [39]. Sample displacement, absorption, and Lorentz-polarization corrections were applied. The crystallinity index (CI) was calculated using the TOPAS v6 built-in tool as the ratio of crystalline area to total area.

### Atomic force microscopy (AFM)

AFM images were collected using a Solver PRO Scanning Probe Microscope (NT-MDT, Netherlands) in semi-contact mode, employing an AFM probe with a silicon cantilever with a force constant of 8-42 N/m and a resonance frequency of 200-360 kHz. Aqueous suspensions of CNC and CNF (0.005 wt%, 20 µL) were placed on newly split mica surfaces and dried in a desiccator overnight. The AFM scan rate was 0.30 Hz, and the image resolution was 512×512 px for 5×5 µm and 20×20 µm. AFM data were processed in Gwyddion [84].

### Particle size analysis

Particle size distributions of α-chitin, β-chitin were measured by laser diffraction on a Mastersizer 3000 Hydro MV (Malvern, UK), using water as dispersant. Refractive indices of 1.468 and 1.330 were used for substrate and water, respectively. The particle absorption index was set to 0.01. Five parallel measurements were run for each substrate. For calculation of specific surface area (SSA), we assumed spherical particles: SSA = 6∕(*pD*[3, 2]), where *ρ* is the particle density (set to 1.43), and *D*[3, 2] is the surface-area-weighted mean diameter.

### Bacterial strains and plasmids

*Sc*CBM2 genes were cloned into the pNIC-CH expression vector (AddGene) carrying a C-terminal Hisx6 tag and conferring kanamycin resistance, as described previously [85]. The plasmid for *Ba*CBM2 was prepared as described previously [14]. The construct was based on the sequence obtained from the C-terminal CBM2 of *B. anthracis* chitinase B, which is an extended version (21 residues more) of the *Ba*CBM2-t variant presented by Weber *et al.* [13] (GenBank: QBK17415.1). The backbone plasmid was pET-21b(+) carrying an N-terminal Hisx6 tag and conferring ampicillin resistance. *E. coli* BL21 (DE3) cells (New England Biolabs, USA, catalog number C2527) were transformed with the *Sc*CBM2 and *Ba*CBM2 plasmids using a standard heat-shock transformation protocol. Plasmids for GFP-*Sc*CBM2 and GFP-*Sc*CBM2-H69W were synthesized by Twist Bioscience (USA) and subcloned into the pET28a(+) vector. The constructs carried an N-terminal Hisx6 tag and a green fluorescent protein (GFP) and conferred kanamycin resistance. E10T point mutations were introduced into the GFP-*Sc*CBM2 and GFP-*Sc*CBM2-H69W constructs by site-directed mutagenesis using the forward primer 5’-CTATAGCGTGaccAACAGCTGGTC-3’ and the reverse primer 5’-ACGGCCATACAACCC-3’. The plasmids were transformed into *E. coli* T7 Express cells (New England Biolabs, catalog number C2566) by heat shock.

### Protein expression and purification

The cells of interest were inoculated into 5 mL LB medium (10 g/L tryptone, 5 g/L yeast extract, 5 g/L NaCl) supplemented with 50 µg/mL kanamycin or 100 µg/mL ampicillin, and incubated at 37^◦^C, 225 rpm overnight. For production of non-isotopically labeled protein, 500 mL 2xLB medium (20 g/L tryptone, 10 g/L yeast extract, 5 g/L NaCl) supplemented with 50 µg/mL kanamycin or 100 µg/mL ampicillin was inoculated with 1% pre-culture. For production of ^15^N-labeled protein, 1% pre-culture was used to inoculate 500 mL M9 media (6 g/L Na_2_HPO_4_, 3 g/L KH_2_PO, 0.5 g/L NaCl) supplemented with 50 µg/mL kanamycin or 100 µg/mL ampicillin, 2 mM MgSO_4_, 0.1 mM CaCl_2_, 10 mL BioExpress Cell Growth Media (Cambridge Isotope Laboratories, USA), 5 mL Gibco^TM^ MEM Vitamin Solution (100x, Thermo Fisher Scientific, USA), 500 µL trace metal solution (1000x, containing 10 g/L CaCl_2_, 8 g/L MnSO_4_, 5 g/L FeSO_4_, 1 g/L CuSO_4_, and 1 g/L ZnSO_4_), 1 g/L (^15^NH_4_)_2_SO_4_ and 1% glycerol. The main cultures were incubated either in a shaking incubator at 22°C and 225 rpm or in a LEX bioreactor (Epiphyte3, Canada) at 22°C. At OD_600_ = 1.6-1.9, protein expression was induced by adding 0.1 mM isopropyl-β-D-1-thiogalactopyranoside (IPTG) followed by incubation for 24 h at 22°C. The cells were harvested by centrifugation at 4°C and 5000*g* for 10 min. The pellet was resuspended in 25 mL of 50 mM Tris-HCl (pH 8.0) supplemented with 300 mM NaCl and one EDTA-free protease inhibitor tablet (Roche). The cells were lysed by sonication for 10 min (5 s on, 10 s off), and the cell debris was separated from the solution by centrifugation at 4°C, 25000*g* for 30 min.

The supernatant was filtered through a 0.45 µm filter prior to purification by Ni affinity chromatography on an Äkta pure FPLC system (Cytiva, USA) using a 1 mL HisTrap HP column (Cytiva). The protein was eluted by applying a 0–400 µM imidazole gradient in 50 mM Tris-HCl (pH 8.0) supplemented with 300 mM NaCl. Pure fractions containing the target protein were identified by SDS-PAGE using 12% SurePAGE Bis-Tris gels (GenScript), with the PAGE-MASTER Protein Standard Plus (GenScript) (***Figure S17***), and concentrated using a Vivaspin® concentrator (Sartorius) with a 5 or 10 kDa cut-off. The buffer was exchanged to the appropriate buffer for each experiment by gel filtration using a PD MidiTrap G-25 column (Cytiva). Protein concentrations were determined by measuring A280 on a NanoDrop ND-1000 spectrophotometer (Thermo Fisher Scientific) using the theoretical extinction coefficient (***Table S1***) calculated using the ProtParam online tool [86].

### Affinity assays

Binding to crystalline PET film was assessed for GFP-labeled *Sc*CBM2 in HEPES buffer (10 mM HEPES supplemented with 150 mM NaCl, 10 % glycerol, pH 7.5), following the protocol described in [14]. Discs of crystalline PET film (0.64 cm^2^ surface area) were cut using an office hole punch and added to fluorescent-suitable black 96-well plates (Greiner, Austria). The discs were washed with 10% SDS for 10 min, followed by deionized water (2×5 min). The discs were incubated in 200 µL HEPES buffer for 30 min. The buffer was removed from the substrates, and 300 µL of protein sample was added. Triplicate samples of concentrations specified in ***Table S2*** were used. The plate was covered with adhesive sheets and incubated at 60°C, 300 rpm for 1 h. 250 µL of each sample was immediately transferred to fresh wells and cooled to rt. Fluorescence intensity was measured on an Infinite 200 PRO plate reader (TECAN, Switzerland) with an excitation wavelength of 450 nm and an emission wavelength of 510 nm.

A modified assay protocol was used to measure binding of GFP-labeled *Sc*CBM2 (wt, E10T, H69W, and E10T-H69W) to the same crystalline PET substrate in phosphate buffer (50 mM sodium phosphate, pH 7.5). Here, the PET discs were washed with 20% ethanol and rinsed twice with deionized water. 300 µL of protein sample was added to Eppendorf tubes. Triplicate (wt) or duplicate (variants) samples of concentrations specified in ***Table S2*** were used. One PET disc was added to each tube, and the tubes were spun down to submerge the discs in the solution. The samples were incubated at rt for 20 min, then at 60°C at 300 rpm for 1 h. 250 µL of each sample was immediately transferred to fluorescent-suitable black 96-well plates (Greiner) and cooled to rt. Fluorescence intensity was measured as described above.

Binding to biopolymer substrates (Avicel, α-chitin, β-chitin, CNF, and CNC) was assessed for non-labeled CBMs by incubating triplicate 200 µL protein samples (concentrations specified in ***Table S2***) in phosphate buffer with 10 g/L biopolymer substrates (5 g/L for CNF and CNC substrates) for 1-2 h at 22°C and 700 rpm. The free protein in solution was then separated from the substrate by filtering the solutions through a 96-well filter plate (Millipore) into a UV-transparent 96-well plate (Corning, USA). 150 µL of each sample was then transferred to neighboring wells, and absorbance at 280 nm was measured on the plate reader.

For all assays, the concentration of free protein [P_free_] was determined using standard curves of each respective protein, where the standards were treated exactly the same as the assay samples, without adding substrate. The concentration of bound protein [P_bound_] was calculated by subtracting the measured free protein concentration [P_free_] from the total protein concentration. Binding parameters were estimated by fitting the binding data to a one-site binding isotherm [P_bound_] = *B*_max_[P_free_]∕(*K_d_* + [P_free_]).

### Nuclear magnetic resonance (NMR)

NMR spectra were recorded on a Bruker Ascend 800 MHz NMR spectrometer (Bruker BioSpin AG, Fälladen, Switzerland) equipped with an Avance NEO console and a 5 mm cryogenic CP-TCI probe at the NV-NMR Center at NTNU, which is part of the Norwegian NMR Platform (NNP). We have deposited the assignment of *Ba*CBM2 in the Biological Magnetic Resonance Data Bank (BMRB: 53987). The NMR assignment of *Sc*CBM2 (BMRB: 27078) has been published previously [85].

NMR spectra were recorded and processed in Bruker TopSpin version 4.4.0. Nuclear relaxation rates (*T*_1_, *T*_2_) were measured as follows: ^15^N-*T*_1_ was measured using a pseudo-3D ^15^N-HSQC experiment (hsqct1etf3gpsi3d.2), with the third dimension representing variable relaxation time delays (0.005, 0.1, 0.2, 0.5, 1, 1.5, 2, 3, and 4 s). ^1^H-*T*_1_ was measured using a series of ^15^N-HSQC experiments modified with a saturation-recovery pulse-sequence block, where the recovery delay was varied between experiments (0.25, 0.5, 1, 2, 4, and 6 s). ^1^H-*T*_2_ was measured using a series of ^15^N-HSQC experiments modified with a Carr-Purcell-Meiboom-Gill (cpmg) pulse sequence block repeated a variable number of times, corresponding to a variable relaxation delay between experiments (1.85, 3.7, 7.4, 11.1, 18.5, and 46.25 ms). For most experiments, water flip-back was applied to minimize solvent signals. Relaxation times and relaxation rates (*R*_1_, *R*_2_) were calculated by fitting peak intensities to the appropriate functions in Bruker Dynamics Center version 2.8.3.

Chemical shift perturbations (CSP, Δ*δ*) were obtained by calculating the shift in amide peak positions in ^15^N-HSQC spectra as 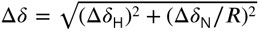, where Δ*δ_H_* and Δ*δ_N_* are the CSP of the amide proton and nitrogen in ppm, and the chemical shift scaling factor *R* was set to 6.5 [87]. This was done using CcpNmr AnalysisAssign [88]. Solvent paramagnetic relaxation enhancement (sPRE) was calculated from measured *R*_1_ and *R*_2_ relaxation rates in the presence of increasing concentrations *c* of gadodiamide (4 points, *c* = 0–1 mM), by linear regression of 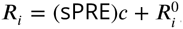, where *i* = 1 or 2, and 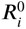 is the relaxation rate in the absence of gadodiamide. Changes in sPRE were calculated as ΔsPRE = sPRE_s_ − sPRE_f_, where the subscripts s and f denote the presence and absence of substrate, respectively. Changes in relaxation rates were calculated as the ratio of the relaxation rate in the presence and absence of substrate, Δ*R_i_* = *R_i,_*_s_∕*R_i,_*_f_.

NMR samples consisted of ^15^N-labeled *Ba*CBM2 (107 µM) in 25 mM NaAc supplemented with 10 mM NaCl and 10% D_2_O (pH 5.0), and ^15^N-labeled *Sc*CBM2 (82 µM) in 50 mM sodium phosphate supplemented with 10% D_2_O (pH 8.0). Samples containing substrate were supplemented with 5 mM bis(2-hydroxyethyl) terephthalate (BHET) or 1.5 mg/mL PET nanoparticles.

### Simulations

Starting structures for *Sc*CBM2, *Sc*CBM2-H69W, *Sc*CBM2-E10T, *Sc*CBM2-E10T-H69W, and *Ba*CBM2 were generated from AlphaFold3 [31] models. Each system was solvated with TIP3P water in a dodecahedral box with a minimum solute-wall distance of 1.0 nm and neutralized with Na^+^ and Cl^-^. All simulations were performed with GROMACS 2022.3 [89] using the Amber99SB-ILDN [90] force field. Each system was energy-minimized using the steepest-descent algorithm for 50000 steps. Equilibration was carried out in two stages: (i) 100 ps in the NVT ensemble at 300 K using the V-rescale thermostat [91] with separate coupling for protein and solvent, and (ii) 100 ps in the NPT ensemble at 300 K and 1 bar using the Nosé-Hoover thermostat [92, 93] and Parrinello-Rahman barostat [94]. Production runs of 250 ns were conducted in the NPT ensemble using a 2 fs time step, with bonds involving hydrogen constrained by the LINCS algorithm [95]. The Nose-Hoover thermostat and Parrinello-Rahman barostat were used with the same settings as the equilibration step. Coordinates were written every 100 ps, yielding 10000 frames per trajectory.

Radial distribution functions (RDFs) between the protein and water oxygen atoms were calculated using the GROMACS rdf tool. To assess variability, each production trajectory was divided into five blocks of equal length, and an RDF was computed independently for each block. The mean RDF and standard error of the mean (SEM) across blocks were then calculated for each CBM. Three-dimensional water density maps were computed from the production trajectories using MDAnalysis [96, 97]. Water oxygen atoms were binned onto a grid with 4.0 Å spacing and a 2.0 Å padding around the solute using the DensityAnalysis module, with all frames aligned to the protein via mass-weighted rotational and translational fitting after unwrapping. The resulting densities were converted to units of bulk water density using the TIP3P conversion factor and exported in OpenDX format. The density map was then loaded into PyMOL [62] and projected onto the van der Waals surface of each CBM using a continuous color ramp, such that surface regions with high local water density (relative to bulk) are highlighted. This approach was used to visualize differences in hydration patterns on the binding surfaces of *Ba*CBM2 and *Sc*CBM2.

## Data availability

Data and scripts used for analysis and for generating the figures are available from https://github.com/gcourtade/papers/blob/master/2026/CBM2/.

## Supporting information

This article contains supporting information.

## Acknowledgments

This work was supported by the Novo Nordisk Foundation (grant number: NNF22OC0073963; to GC) and the Independent Research Fund Denmark | Technology and Production Sciences (grant number: 1032-00273B; to MSM). We are grateful to Dr. Ellinor B. Heggset (RISE PFI AS) for providing us with CNC and CNF substrates.

## Notes

https://github.com/gcourtade/papers/blob/master/2026/CBM2/

### Competing Interest Statement

The authors have declared no competing interest.

## References

[1] A Boraston, D Bolam, H Gilbert, G Davies, Carbohydrate-binding modules: Fine-tuning polysaccharide recognition. Biochemical Journal 382, 769–781 (2004).

[2] C Hervé, et al., Carbohydrate-binding modules promote the enzymatic deconstruction of intact plant cell walls by targeting and proximity effects. Proceedings of the National Academy of Sciences 107, 15293–15298 (2010).

[3] D Chettri, AK Verma, M Chirania, AK Verma, Unlocking the potential of carbohydrate-binding modules: Functional insights, engineering strategies, and applications. Cellulose 33, 4369–4400 (2026).

[4] P Tomme, et al., Characterization and affinity applications of cellulose-binding domains. Journal of Chromatography B: Biomedical Sciences and Applications 715, 283–296 (1998).

[5] PJ Simpson, H Xie, DN Bolam, HJ Gilbert, MP Williamson, The structural basis for the ligand specificity of family 2 carbohydrate-binding modules. Journal of Biological Chemistry 275, 41137–41142 (2000).

[6] AW Blake, et al., Understanding the biological rationale for the diversity of cellulose-directed carbohydrate-binding modules in prokaryotic enzymes. Journal of Biological Chemistry 281, 29321–29329 (2006).

[7] J Lehtiö, et al., The binding specificity and affinity determinants of family 1 and family 3 cellulose binding modules. Proceedings of the National Academy of Sciences 100, 484–489 (2003).

[8] J Guo, JM Catchmark, Binding specificity and thermodynamics of cellulose-binding modules from *Trichoderma reesei* Cel7A and Cel6A. Biomacromolecules 14, 1268–1277 (2013).

[9] S Arola, MB Linder, Binding of cellulose binding modules reveal differences between cellulose substrates. Scientific Reports 6, 35358 (2016).

[10] C Yang, et al., Interactions between the engineering family 3 carbohydrate-binding module (CBM3) and different types of nanocelluloses revealed by single-molecule force spectroscopy (SMFS). International Journal of Biological Macromolecules 352, 151159 (2026).

[11] A Creagh, E Ong, E Jervis, D Kilburn, C Haynes, Binding of the cellulose-binding domain of exoglucanase Cex from *Cellulomonas fimi* to insoluble microcrystalline cellulose is entropically driven. Proceedings of the National Academy of Sciences 93, 12229–34 (1996).

[12] BW McLean, et al., Carbohydrate-binding modules recognize fine substructures of cellulose. Journal of Biological Chemistry 277, 50245–50254 (2002).

[13] J Weber, et al., Interaction of carbohydrate-binding modules with poly(ethylene terephthalate). Applied Microbiology and Biotechnology 103, 4801–4812 (2019).

[14] AP Rennison, P Westh, MS Møller, Protein-plastic interactions: The driving forces behind the high affinity of a carbohydrate-binding module for polyethylene terephthalate. Science of The Total Environment 870, 161948 (2023).

[15] Y Zhang, L Wang, J Chen, J Wu, Enhanced activity toward PET by site-directed mutagenesis of *Thermobifida fusca* cutinase–CBM fusion protein. Carbohydrate Polymers 97, 124–129 (2013).

[16] L Dai, et al., Enhancing PET hydrolytic enzyme activity by fusion of the cellulose–binding domain of cellobiohydrolase I from *Trichoderma reesei*. Journal of Biotechnology 334, 47–50 (2021).

[17] R Graham, et al., The role of binding modules in enzymatic poly(ethylene terephthalate) hydrolysis at high-solids loadings. Chem Catalysis 2, 2644–2657 (2022).

[18] Y Chen, et al., Construction of fusion protein with carbohydrate-binding module and leaf-branch compost cutinase to enhance the degradation efficiency of polyethylene terephthalate. International Journal of Molecular Sciences 24, 2780 (2023).

[19] AP Rennison, A Prestel, P Westh, MS Møller, Comparative biochemistry of PET hydrolase-carbohydrate-binding module fusion enzymes on a variety of PET substrates. Enzyme and Microbial Technology 180, 110479 (2024).

[20] Y Liu, et al., Enhanced degradation activity of PET plastics by fusion protein of anchor peptide LCI and *Thermobifida fusca* cutinase. Enzyme and Microbial Technology 184, 110562 (2025).

[21] AP Rennison, et al., A high-throughput investigation of the binding specificity of carbohydrate-binding modules for synthetic and natural polymers. ChemSusChem 18, e202500468 (2025).

[22] AA Bogan, KS Thorn, Anatomy of hot spots in protein interfaces. Journal of Molecular Biology 280, 1–9 (1998).

[23] P Tomme, AL Creagh, DG Kilburn, CA Haynes, Interaction of polysaccharides with the N-terminal cellulose-binding domain of *Cellu*lomonas fimi CenC. 1. Binding specificity and calorimetric analysis. Biochemistry 35, 13885–13894 (1996).

[24] Y Zhang, et al., Interactions between type A carbohydrate binding modules and cellulose studied with a quartz crystal microbalance with dissipation monitoring. Cellulose 27, 3661–3675 (2020).

[25] A Boraston, The interaction of carbohydrate-binding modules with insoluble non-crystalline cellulose is enthalpically driven. Biochemical Journal 385, 479–484 (2005).

[26] T Liu, et al., Binding affinity of family 4 carbohydrate binding module on cellulose films of nanocrystals and nanofibrils. Carbohydrate Polymers 251, 116725 (2021).

[27] Y You, et al., Carbohydrate binding modules: Compact yet potent accessories in the specific substrate binding and performance evolution of carbohydrate-active enzymes. Biotechnology Advances 73, 108365 (2024).

[28] S Li, G Liu, Harnessing cellulose-binding protein domains for the development of functionalized cellulose materials. Bioresources and Bioprocessing 11, 74 (2024).

[29] F Madeira, et al., The EMBL-EBI Job Dispatcher sequence analysis tools framework in 2024. Nucleic Acids Res 52, W521–W525 (2024).

[30] GE Crooks, G Hon, JM Chandonia, SE Brenner, WebLogo: A sequence logo generator. Genome Research 14, 1188–1190 (2004).

[31] J Abramson, et al., Accurate structure prediction of biomolecular interactions with AlphaFold 3. Nature 630, 493–500 (2024).

[32] AM Waterhouse, JB Procter, DMA Martin, M Clamp, GJ Barton, Jalview Version 2—a multiple sequence alignment editor and analysis workbench. Bioinformatics 25, 1189–1191 (2009).

[33] L Segal, J Creely, A MartinJr, C Conrad, An empirical method for estimating the degree of crystallinity of native cellulose using the X-ray diffractometer. Textile Research Journal 29, 786–794 (1959).

[34] KS Salem, et al., Comparison and assessment of methods for cellulose crystallinity determination. Chemical Society Reviews 52, 6417–6446 (2023).

[35] AD French, Increment in evolution of cellulose crystallinity analysis. Cellulose 27, 5445–5448 (2020).

[36] P Ahvenainen, I Kontro, K Svedström, Comparison of sample crystallinity determination methods by X-ray diffraction for challenging cellulose I materials. Cellulose 23, 1073–1086 (2016).

[37] Y Nishiyama, P Langan, H Chanzy, Crystal structure and hydrogen-bonding system in cellulose Iβ from synchrotron X-ray and neutron fiber diffraction. Journal of the American Chemical Society 124, 9074–9082 (2002).

[38] JA Kaduk, PDF Card 00-056-1717. Private communication (2005).

[39] X Ju, M Bowden, EE Brown, X Zhang, An improved X-ray diffraction method for cellulose crystallinity measurement. Carbohydrate Polymers 123, 476–481 (2015).

[40] J Li, et al., Effects of hydrolysis conditions on the morphology of cellulose II nanocrystals (CNC-II) derived from mercerized microcrystalline cellulose. International Journal of Biological Macromolecules 258, 128936 (2024).

[41] A Abbasi Moud, Cellulose nanocrystals examined by atomic force microscopy: Applications and fundamentals. ACS Food Science & Technology 2, 1789–1818 (2022).

[42] Y Qing, et al., A comparative study of cellulose nanofibrils disintegrated via multiple processing approaches. Carbohydrate Polymers 97, 226–234 (2013).

[43] P Lekha, et al., Effect of mechanical treatment on properties of cellulose nanofibrils produced from bleached hardwood and softwood pulps. Maderas. Ciencia y tecnología 18, 457–466 (2016).

[44] ML Di Lorenzo, Crystallization of poly(ethylene terephthalate): A review. Polymers 16, 1975 (2024).

[45] M Hirai, et al., Direct evidence for the effect of glycerol on protein hydration and thermal structural transition. Biophysical Journal 115, 313–327 (2018).

[46] M Wang, et al., The role of glycerol in preserving proteins needs to be reconsidered. ACS Sustainable Chemistry & Engineering 10, 15175–15185 (2022).

[47] BJM Rooijakkers, MS Ikonen, MB Linder, Fungal-type carbohydrate binding modules from the coccol-ithophore *Emiliania huxleyi* show binding affinity to cellulose and chitin. PLOS ONE 13, e0197875 (2018).

[48] Y Yamauchi, PY Apel, Adsorption of anionic surfactant on porous and non-porous polyethylene terephthalate films. Colloid Journal 79, 286–294 (2017).

[49] M Furukawa, N Kawakami, K Oda, K Miyamoto, Acceleration of enzymatic degradation of poly(ethylene terephthalate) by surface coating with anionic surfactants. ChemSusChem 11, 4018–4025 (2018).

[50] M Furukawa, N Kawakami, A Tomizawa, K Miyamoto, Efficient degradation of poly(ethylene terephthalate) with *Thermobifida fusca* cutinase exhibiting improved catalytic activity generated using mutagenesis and additive-based approaches. Scientific Reports 9, 16038 (2019).

[51] G Vaaje-Kolstad, SJ Horn, DM van Aalten, B Synstad, VG Eijsink, The non-catalytic chitin-binding protein CBP21 from serratia marcescens is essential for chitin degradation. Journal of Biological Chemistry 280, 28492–28497 (2005).

[52] K Manjeet, P Purushotham, C Neeraja, AR Podile, Bacterial chitin binding proteins show differential substrate binding and synergy with chitinases. Microbiological Research 168, 461–468 (2013).

[53] G Courtade, Z Forsberg, EB Heggset, VG Eijsink, FL Aachmann, The carbohydrate-binding module and linker of a modular lytic polysaccharide monooxygenase promote localized cellulose oxidation. Journal of Biological Chemistry 293, 13006–13015 (2018).

[54] GT Beckham, et al., Identification of amino acids responsible for processivity in a family 1 carbohydrate-binding module from a fungal cellulase. The Journal of Physical Chemistry B 114, 1447–1453 (2010).

[55] DJ Dagel, et al., In situ imaging of single carbohydrate-binding modules on cellulose microfibrils. The Journal of Physical Chemistry B 115, 635–641 (2011).

[56] J Široký, et al., Analysis of crystallinity changes in cellulose II polymers using carbohydrate-binding modules. Carbohydrate Polymers 89, 213–221 (2012).

[57] M Sahihi, et al., Probing enzymatic PET degradation: molecular dynamics analysis of cutinase adsorption and stability. Journal of Chemical Information and Modeling 64, 4112–4120 (2024).

[58] DA Torchia, Dynamics of biomolecules from picoseconds to seconds at atomic resolution. Journal of Magnetic Resonance 212, 1–10 (2011).

[59] C Charlier, SF Cousin, F Ferrage, Protein dynamics from nuclear magnetic relaxation. Chemical Society Reviews 45, 2410–2422 (2016).

[60] G Pintacuda, G Otting, Identification of protein surfaces by NMR measurements with a paramagnetic Gd(III) chelate. Journal of the American Chemical Society 124, 372–373 (2002).

[61] AJ Lenard, FA Mulder, T Madl, Solvent paramagnetic relaxation enhancement as a versatile method for studying structure and dynamics of biomolecular systems. Progress in Nuclear Magnetic Resonance Spectroscopy 132-133, 113–139 (2022).

[62] The PyMOL Molecular Graphics System, Version 3.1.5.1 (2025).

[63] SF Badino, JA Bååth, K Borch, K Jensen, P Westh, Adsorption of enzymes with hydrolytic activity on polyethylene terephthalate. Enzyme and Microbial Technology 152, 109937 (2021).

[64] A Jäckering, et al., From bulk to binding: Decoding the entry of PET into hydrolase binding pockets. JACS Au 4, 4000–4012 (2024).

[65] A Nakamura, N Kobayashi, N Koga, R Iino, Positive charge introduction on the surface of thermostabilized PET hydrolase facilitates PET binding and degradation. ACS Catalysis 11, 8550–8564 (2021).

[66] HY Sagong, et al., Implications for the PET decomposition mechanism through similarity and dissimilarity between PETases from *Rhizobacter gummiphilus* and *Ideonella sakaiensis*. Journal of Hazardous Materials 416, 126075 (2021).

[67] Z Kou, D Tolmachev, M Vuorte, M Sammalkorpi, Interactions of NaCl with cellulose Iβ crystal surfaces and the effect on cellulose hydration: A molecular dynamics study. Cellulose 31, 4115–4129 (2024).

[68] Y Fu, WR Busing, Y Jin, KA Affholter, B Wunderlich, Poly(ethylene terephthalate) fibers. 1. Crystal structure and morphology studies with full-pattern x-ray diffraction refinement. Macromolecules 26, 2187–2193 (1993).

[69] Tormo J, et al., Crystal structure of a bacterial family-III cellulose-binding domain: A general mechanism for attachment to cellulose. The EMBO Journal 15, 5739–5751 (1996).

[70] JR King, CM Bowers, EJ Toone, Specific binding at the cellulose binding module–cellulose interface observed by force spectroscopy. Langmuir 31, 3431–3440 (2015).

[71] AM Keys, DW Kastner, LL Kiessling, HJ Kulik, CH-interactions confer orientational flexibility in protein–carbohydrate binding sites. Journal of Biological Chemistry 301, 110379 (2025).

[72] V Spiwok, CH/π interactions in carbohydrate recognition. Molecules 22, 1038 (2017).

[73] N Georgelis, NH Yennawar, DJ Cosgrove, Structural basis for entropy-driven cellulose binding by a type-A cellulose-binding module (CBM) and bacterial expansin. Proceedings of the National Academy of Sciences 109, 14830–14835 (2012).

[74] AP Rennison, A Nousi, P Westh, R Marie, MS Møller, Unveiling PET hydrolase surface dynamics through fluorescence microscopy. ChemBioChem 25, e202300661 (2024).

[75] T Wang, et al., Molecular engineering of PETase for efficient PET biodegradation. Ecotoxicology and Environmental Safety 280, 116540 (2024).

[76] GM Clore, Exploring sparsely populated states of macromolecules by diamagnetic and paramagnetic NMR relaxation. Protein Science 20, 229–246 (2011).

[77] MP Williamson, Using chemical shift perturbation to characterise ligand binding. Progress in Nuclear Magnetic Resonance Spectroscopy 73, 1–16 (2013).

[78] NJ Anthis, GM Clore, Visualizing transient dark states by NMR spectroscopy. Quarterly Reviews of Biophysics 48, 35–116 (2015).

[79] TK Egner, P Naik, NC Nelson, II Slowing, V Venditti, Mechanistic insight into nanoparticle surface adsorption by solution NMR spectroscopy in an aqueous gel. Angewandte Chemie International Edition 56, 9802–9806 (2017).

[80] A Ceccon, V Tugarinov, GM Clore, TiO_2_ nanoparticles catalyze oxidation of huntingtin exon 1-derived peptides impeding aggregation: a quantitative NMR study of binding and kinetics. Journal of the American Chemical Society 141, 94–97 (2018).

[81] G Jameson, X Xiang, R Brüschweiler, Quantitative multistate binding model of silica nanoparticle–protein interactions obtained from multinuclear spin relaxation. The Journal of Physical Chemistry B 126, 9089–9094 (2022).

[82] L Pfaff, D Breite, CP Badenhorst, UT Bornscheuer, R Wei, Chapter Twelve - Fluorimetric high-throughput screening method for polyester hydrolase activity using polyethylene terephthalate nanoparticles in Methods in Enzymology, eds. G Weber, UT Bornscheuer, R Wei. (Academic Press) Vol. 648, pp. 253–270 (2021).

[83] EB Heggset, G Chinga-Carrasco, K Syverud, Temperature stability of nanocellulose dispersions. Carbohydrate Polymers 157, 114–121 (2017).

[84] D Nečas, P Klapetek, Gwyddion: An open-source software for SPM data analysis. Open Physics 10, 181–188 (2012).

[85] G Courtade, Z Forsberg, G Vaaje-Kolstad, VGH Eijsink, FL Aachmann, Chemical shift assignments for the apo-form of the catalytic domain, the linker region, and the carbohydrate-binding domain of the cellulose-active lytic polysaccharide monooxygenase ScLPMO10C. Biomolecular NMR Assignments 11, 257–264 (2017).

[86] E Gasteiger, et al., Protein identification and analysis tools on the ExPASy server in The Proteomics Protocols Handbook, ed. JM Walker. (Humana Press, Totowa, NJ), pp. 571–607 (2005).

[87] FA Mulder, D Schipper, R Bott, R Boelens, Altered flexibility in the substrate-binding site of related native and engineered high-alkaline *Bacillus subtilis*. Journal of Molecular Biology 292, 111–123 (1999).

[88] SP Skinner, et al., CcpNmr AnalysisAssign: A flexible platform for integrated NMR analysis. Journal of Biomolecular NMR 66, 111–124 (2016).

[89] MJ Abraham, et al., GROMACS: High performance molecular simulations through multi-level parallelism from laptops to supercomputers. SoftwareX 1-2, 19–25 (2015).

[90] K Lindorff-Larsen, et al., Improved side-chain torsion potentials for the Amber ff99SB protein force field. Proteins: Structure, Function, and Bioinformatics 78, 1950–1958 (2010).

[91] G Bussi, D Donadio, M Parrinello, Canonical sampling through velocity rescaling. The Journal of Chemical Physics 126, 014101 (2007).

[92] S Nosé, A unified formulation of the constant temperature molecular dynamics methods. Journal of Chemical Physics 81, 511–519 (1984).

[93] W Hoover, Canonical dynamics: Equilibrium phase-space distributions. Physical Review A 31, 1695–1697 (1985).

[94] M Parrinello, A Rahman, Polymorphic transitions in single crystals: A new molecular dynamics method. Journal of Applied Physics 52, 7182–7190 (1981).

[95] B Hess, H Bekker, HJC Berendsen, JGEM Fraaije, LINCS: A linear constraint solver for molecular simulations. Journal of Computational Chemistry 18, 1463–1472 (1997).

[96] N Michaud-Agrawal, EJ Denning, TB Woolf, O Beckstein, MDAnalysis: A toolkit for the analysis of molecular dynamics simulations. Journal of Computational Chemistry 32, 2319–2327 (2011).

[97] RJ Gowers, et al., MDAnalysis: A Python package for the rapid analysis of molecular dynamics simulations in Proceedings of the 15th Python in Science Conference. pp. 98–105 (2016).

