## Supporting information for "Decoding CBM2 binding modes: how similar folds achieve distinct substrate preferences"

[1]Norwegian University of Science and Technology (NTNU), NOBIPOL, Department of Biotechnology and Food Science, Sem Sælands vei 6/8, 7491 Trondheim, Norway
[2]Technical University of Denmark (DTU), Department of Biotechnology and Biomedicine, Søltofts Plads 221/056, 2800 Kongens Lyngby, Denmark
[3]Aalborg University (AAU), Department of Materials and Production, Fibigerstræde 16, 9220 Aalborg, Denmark

**Table S1.** Molecular weight and molar extinction coefficients for expressed proteins.

| Protein | Molecular weight (Da) | Extinction coefficient, $\varepsilon$ ($M^{-1}$ $cm^{-1}$) |
|---|---|---|
| *Sc*CBM2 | 11683 | 23615 |
| *Ba*CBM2 | 10933 | 26470 |
| GFP-*Sc*CBM2 | 39317 | 47120 |
| GFP-*Sc*CBM2-H69W | 39366 | 52620 |
| GFP-*Sc*CBM2-E10T | 39289 | 47120 |
| GFP-*Sc*CBM2-E10T-H69W | 39338 | 52620 |

**Table S2.** Protein concentrations used for affinity assays.

| Assay | Protein concentrations |
|---|---|
| PET | 0, 50, 100, 200, 300, 400, 500, 700, 1000 nM |
| Avicel + ScCBM2 | 0, 2.0, 4.0, 5.6, 8.0, 11.5, 16.0, 21.0 μM |
| Avicel + BaCBM2 | 0, 1.8, 3.6, 6.3, 9.0, 13.5, 27.0, 45.0 μM |
| α-chitin | 0, 1.8, 3.6, 6.3, 9.0, 13.5, 18.0, 27.0, 36.0, 45.0 μM |
| β-chitin | 0, 1.6, 3.2, 5.6, 8.0, 12.0, 16.0, 20.0, 28.0, 36.0 μM |

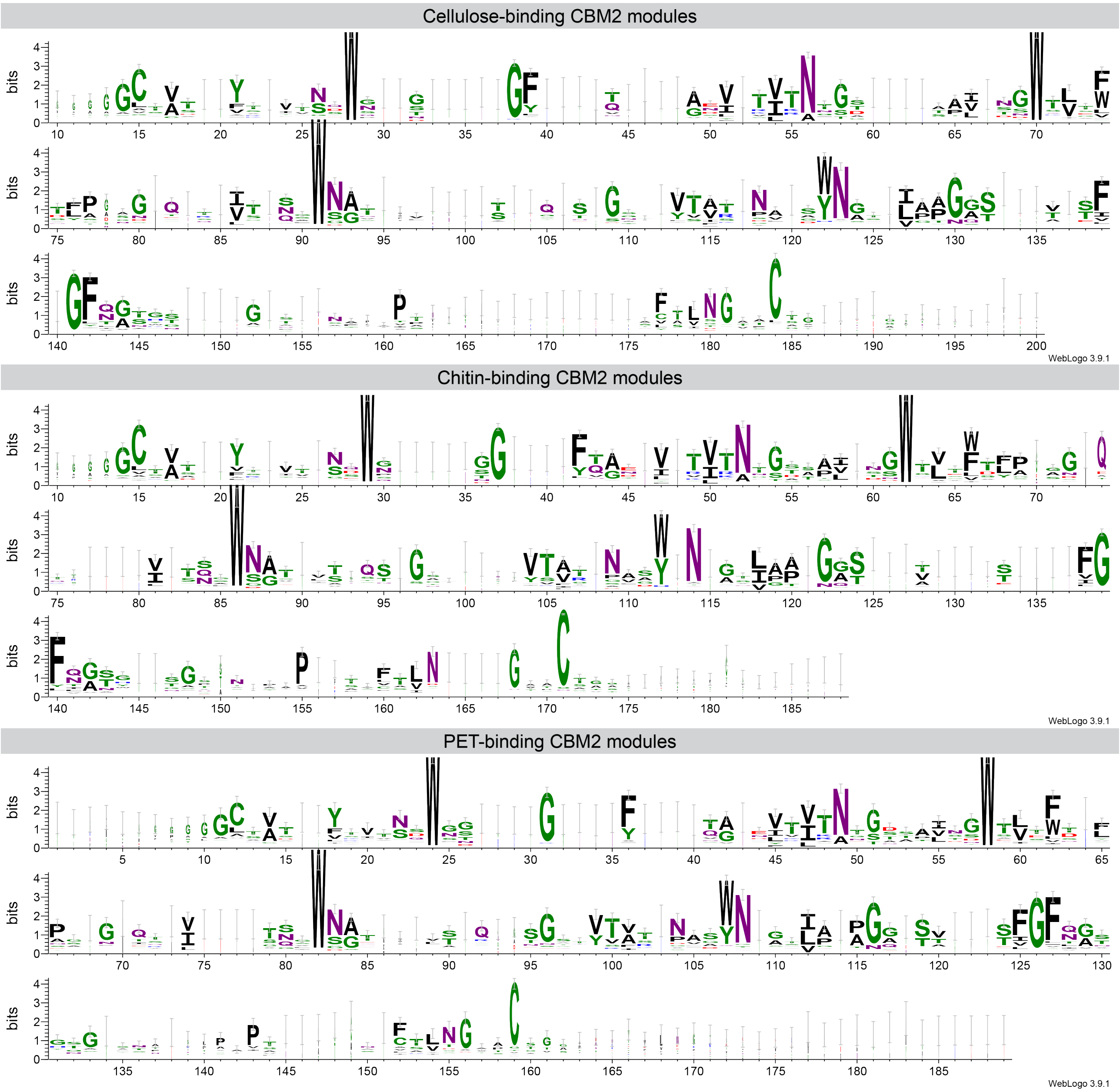


**Figure S1.** Sequence logos for family 2 CBMs classified as cellulose-, chitin- and PET-binders according to Rennison *et al.* [1]. The data set contained 324 cellulose-binders, 322 chitin-binders and 155 PET-binders. The sequences were aligned in Clustal Omega [2], and sequence logos were created using WebLogo 3 [3]. The coloring represents the chemical properties of residues: polar = green, neutral = purple, basic = blue, acidic = red, and hydrophobic = black.

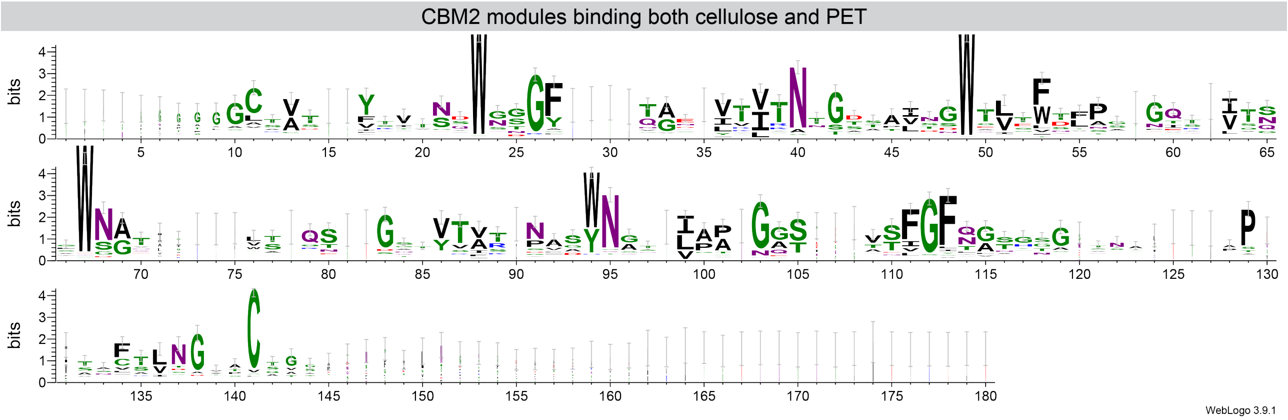


**Figure S2.** Sequence logos for family 2 CBMs classified as both cellulose- and PET-binders according to Rennison *et al.* [1]. The data set contained 123 CBM2s binding to both substrates. The sequences were aligned in Clustal Omega [2], and sequence logos were created using WebLogo 3 [3]. The coloring represents the chemical properties of residues: polar = green, neutral = purple, basic = blue, acidic = red, and hydrophobic = black.

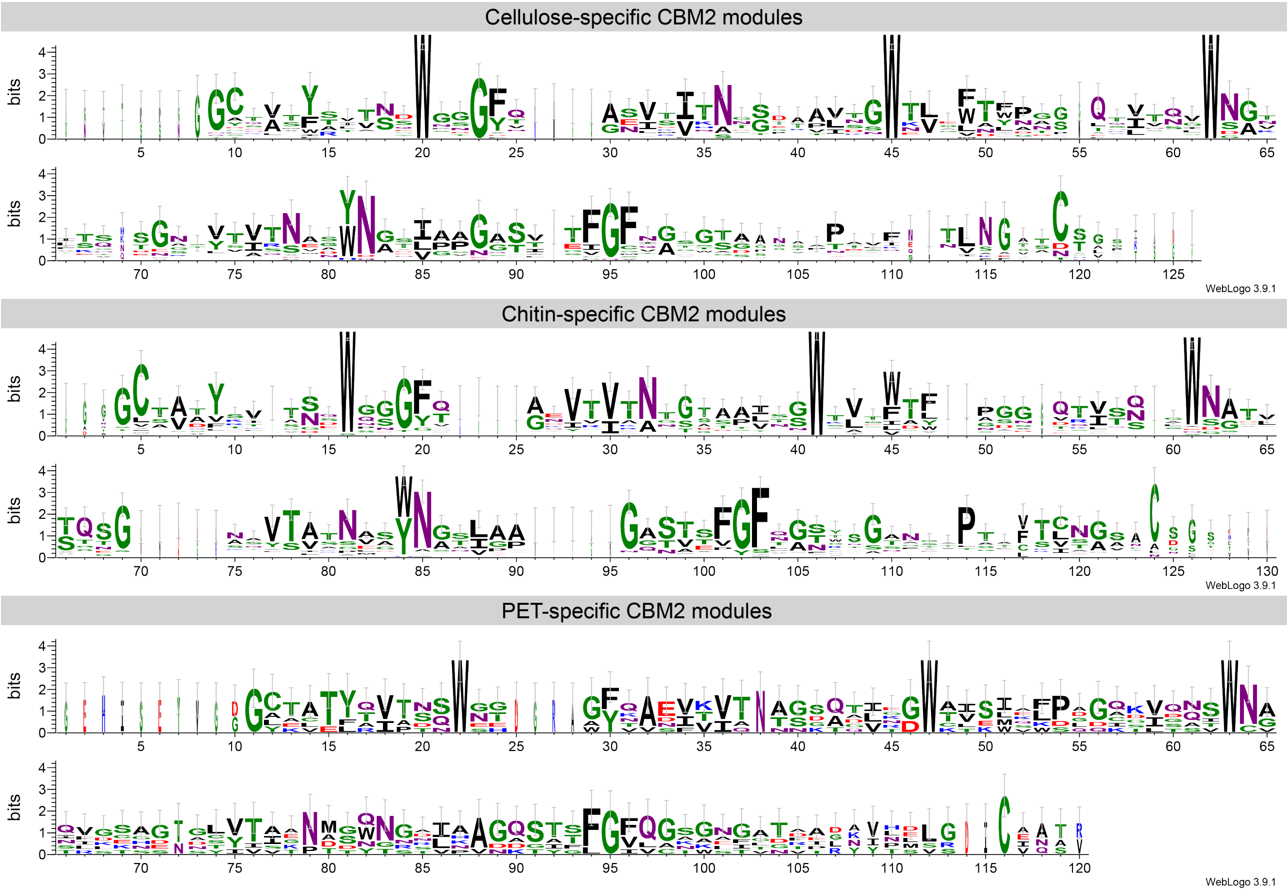


**Figure S3.** Sequence logos for family 2 CBMs classified as highly specific cellulose-, chitin- and PET-binders according to Rennison *et al.* [1]. The data set contained 21 cellulose-specific, 37 chitin-specific and 6 PET-specific CBMs. The sequences were aligned in Clustal Omega [2], and sequence logos were created using WebLogo 3 [3]. The coloring represents the chemical properties of residues: polar = green, neutral = purple, basic = blue, acidic = red, and hydrophobic = black.

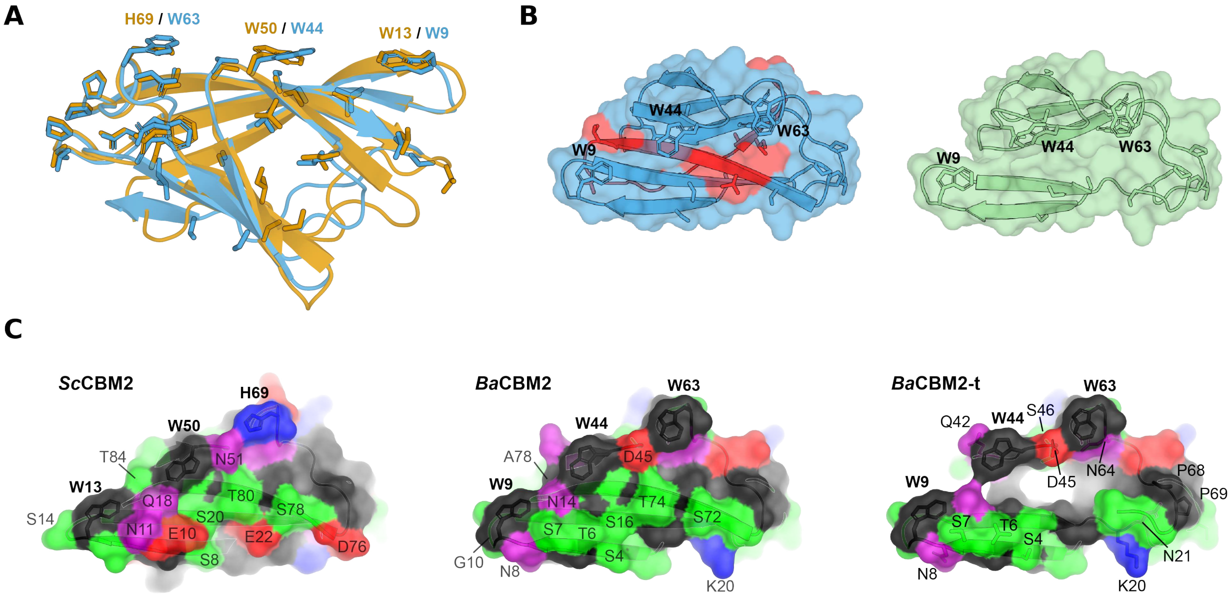


**Figure S4.** AlphaFold [4] structures of *Sc*CBM2 (orange) and *Ba*CBM2 (blue) structurally aligned in PyMOL [5] (RMSD = 0.694) (A). Conserved residues based on amino acid sequence are shown as sticks, and the aromatic triad of the binding surfaces are shown as sticks and labeled with residue codes. AlphaFold structures of *Ba*CBM2 (blue) and *Ba*CBM2-t (green) (B). The 22 residues of the C-terminal of *Ba*CBM2 that are missing in *Ba*CBM2-t are shown in red. AlphaFold structures of *Sc*CBM2, *Ba*CBM2, and *Ba*CBM2-t (C), with residues colored by their chemical properties: polar = green, neutral = purple, basic = blue, acidic = red, and hydrophobic = black. Residues of *Ba*CBM2-t that were shown to interact with PET by Weber *et al.* [6] are shown as sticks and labeled with residue codes.

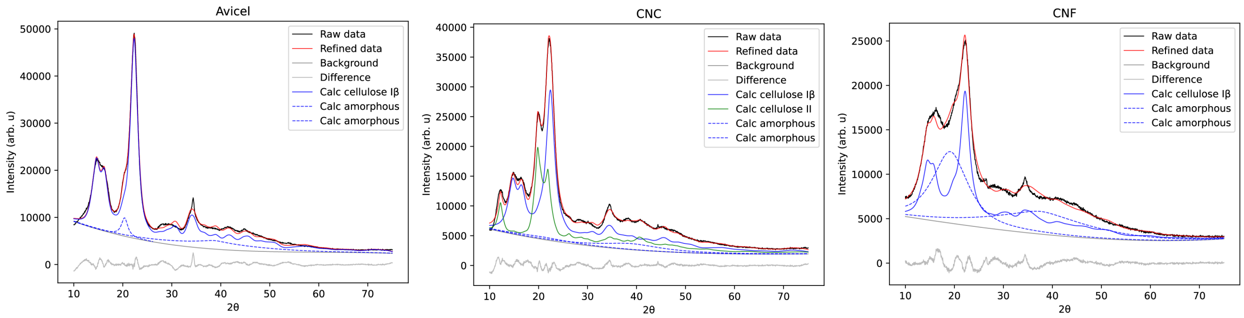


**Figure S5.** XRD patterns of cellulose substrates: Avicel (left), nanocrystalline cellulose (CNC, middle) and cellulose nanofibrils (CNF, right).

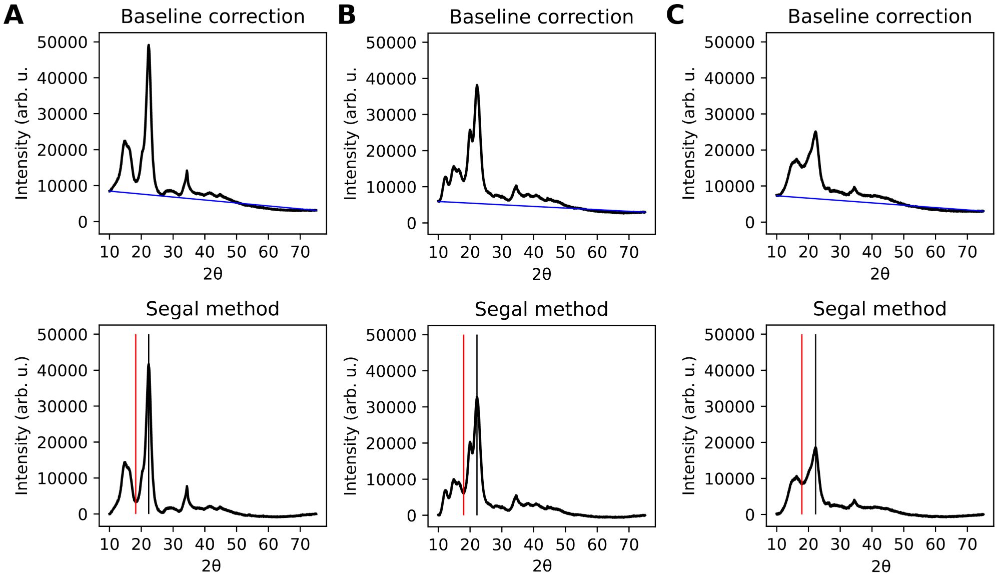


**Figure S6.** Segal method for crystallinity determination of cellulose substrates: Avicel (A), CNC (B) and CNF (C). Baseline correction was performed by subtracting a linear baseline (blue line) from the XRD patterns (top). The maximum intensity $I_C$ of the (2 0 0) peak (indicated by a black line) was used to represent the crystalline fraction, and the minimum intensity $I_A$ around 18° 2θ (indicated by a red line) was used to represent the amorphous fraction. The crystallinity index from the Segal method was calculated as the ratio $(I_C - I_A)/I_C$, which was 92% for Avicel, 82% for CNC and 55% for CNF.

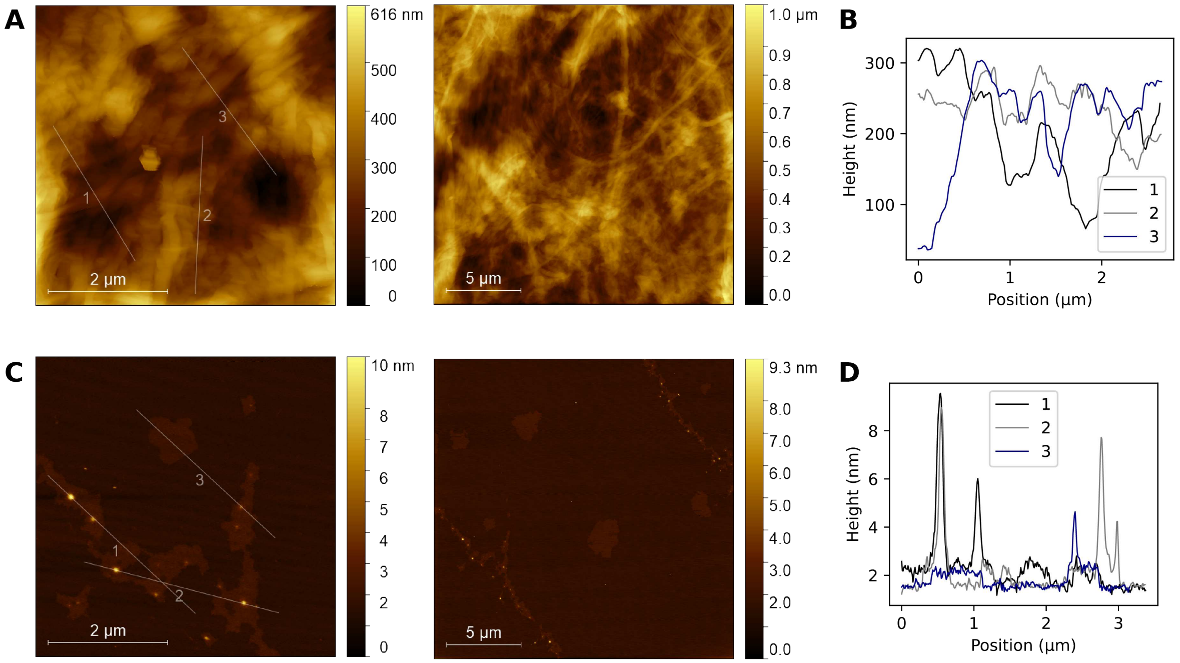


**Figure S7.** AFM images of cellulose nanofibrils (A) and cellulose nanocrystals (C), with scan areas 5×5 µm (left) and 20×20 µm (right). AFM height profiles of CNF (B) and CNC (D), along the lines indicated on the 5×5 µm images.

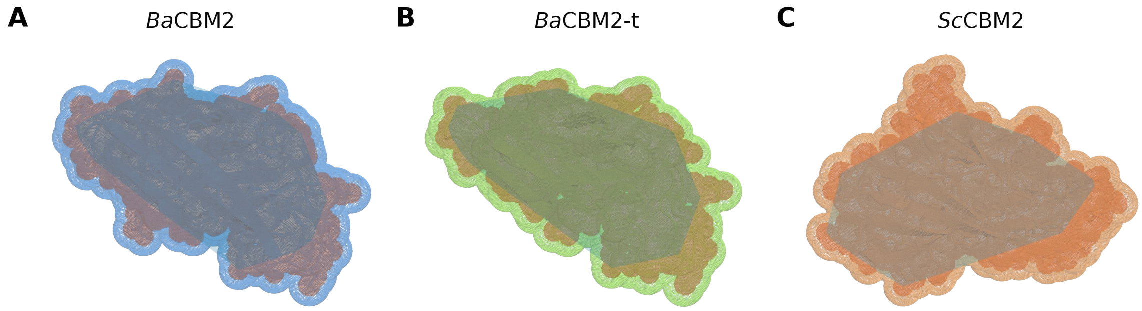


**D**

| | SASA (nm$^2$) | Footprint area (nm$^2$) | Binding patch area (nm$^2$) |
|---|---|---|---|
| BaCBM2 | 51.3 | 8.74 | 6.53 |
| BaCBM2-t | 52.0 | 8.33 | 6.43 |
| ScCBM2 | 55.1 | 9.11 | 6.65 |

**Figure S8.** Protein surface areas and binding site areas for *Ba*CBM2 (A), *Ba*CBM2-t (B), and *Sc*CBM2 (C). The total solvent-accessible surface area (SASA) was calculated using the `get_area` function in PyMol [5] on the AlphaFold [4] structures. We represented the suggested flat binding surfaces on each protein by drawing a plane through three points on the flat surface. We then calculated the total footprint area (red area in A-C) from the projection of all protein atoms (using their van der Waals radius) onto this plane. Additionally, we estimated the area of the flat binding patch by selecting all surface-exposed atoms within 4 Å of the binding plane, projecting the points onto the plane, and generating a convex hull (dark area in A-C) enclosing all points. (D) Calculated SASA, footprint area and binding patch areas.

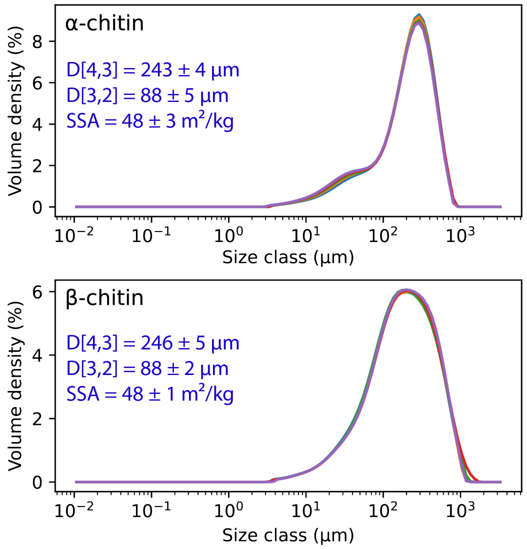


**Figure S9.** Particle size distribution of chitin substrates determined by laser diffraction analysis. The graphs show the distribution of weight classes from 5 identical measurements. Average volume-weighted mean diameter (D[4,5]), average surface-area-weighted mean diameter (D[3,2]), and specific surface area (SSA), assuming spherical particles, are shown for each substrate, with errors representing the standard deviation from the 5 measurements.

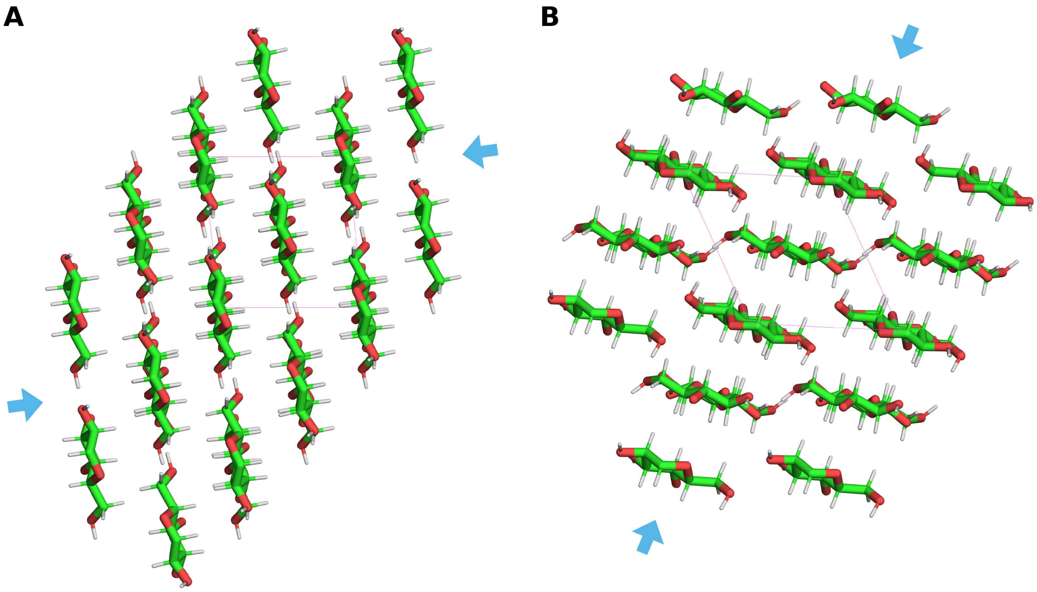


**Figure S10.** Cellulose crystal lattices viewed onto the ab plane of the unit cell (shown in magenta) for cellulose Iβ (A) [7] and cellulose II (B) [8], showing the disadvantageous arrangement of the hydrophobic face of cellulose II. Hydrophobic faces are indicated with blue arrows.

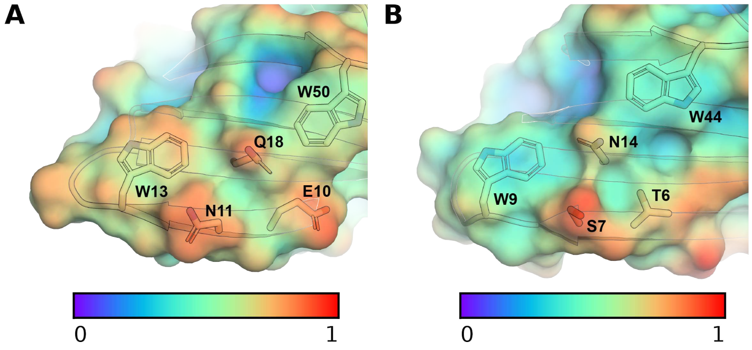


**Figure S11.** Surface water density around key residues around E10 of *Sc*CBM2 (left) and T6 of *Ba*CBM2 (right).

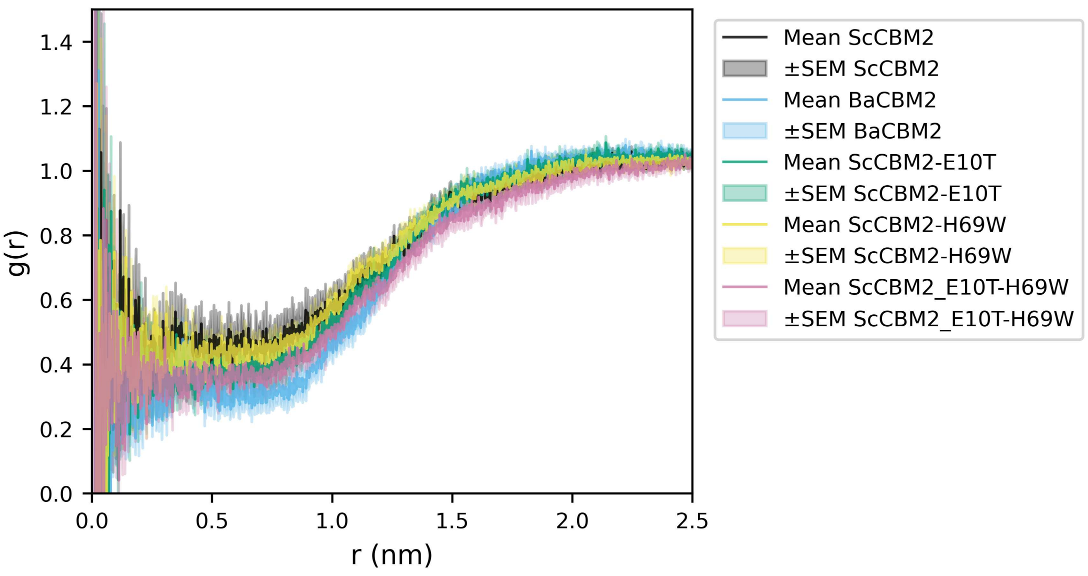


**Figure S12.** Radial distribution functions (RDF) showing the density of water molecules $g(r)$ as a function of distance $r$ from the protein's center of mass during the 250 ns simulation of *Ba*CBM2 and *Sc*CBM2 (wild type and variants E10T, H69W, and E10T-H69W). The error is displayed as the standard error of the mean (SEM) between the calculated RDF from 5 × 50 ns blocks during the simulation.

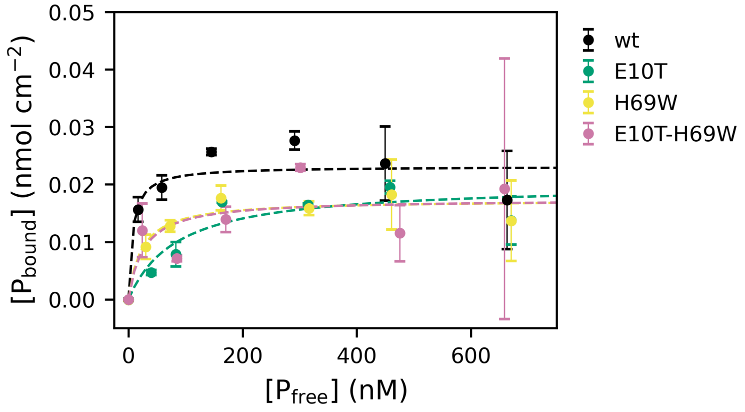


**Figure S13.** Binding of GFP-*Sc*CBM2 (wt, E10T, H69W and E10T-H69W) to crystalline PET film at 60°C. The binding was determined by measuring fluorescence intensity after incubating proteins in 50 mM sodium phosphate at pH 7.5 with PET discs for 1 h. Dissociation constants $K_d$ and binding capacities $B_{max}$ were estimated by fitting the data to a Langmuir isotherm (dashed lines).

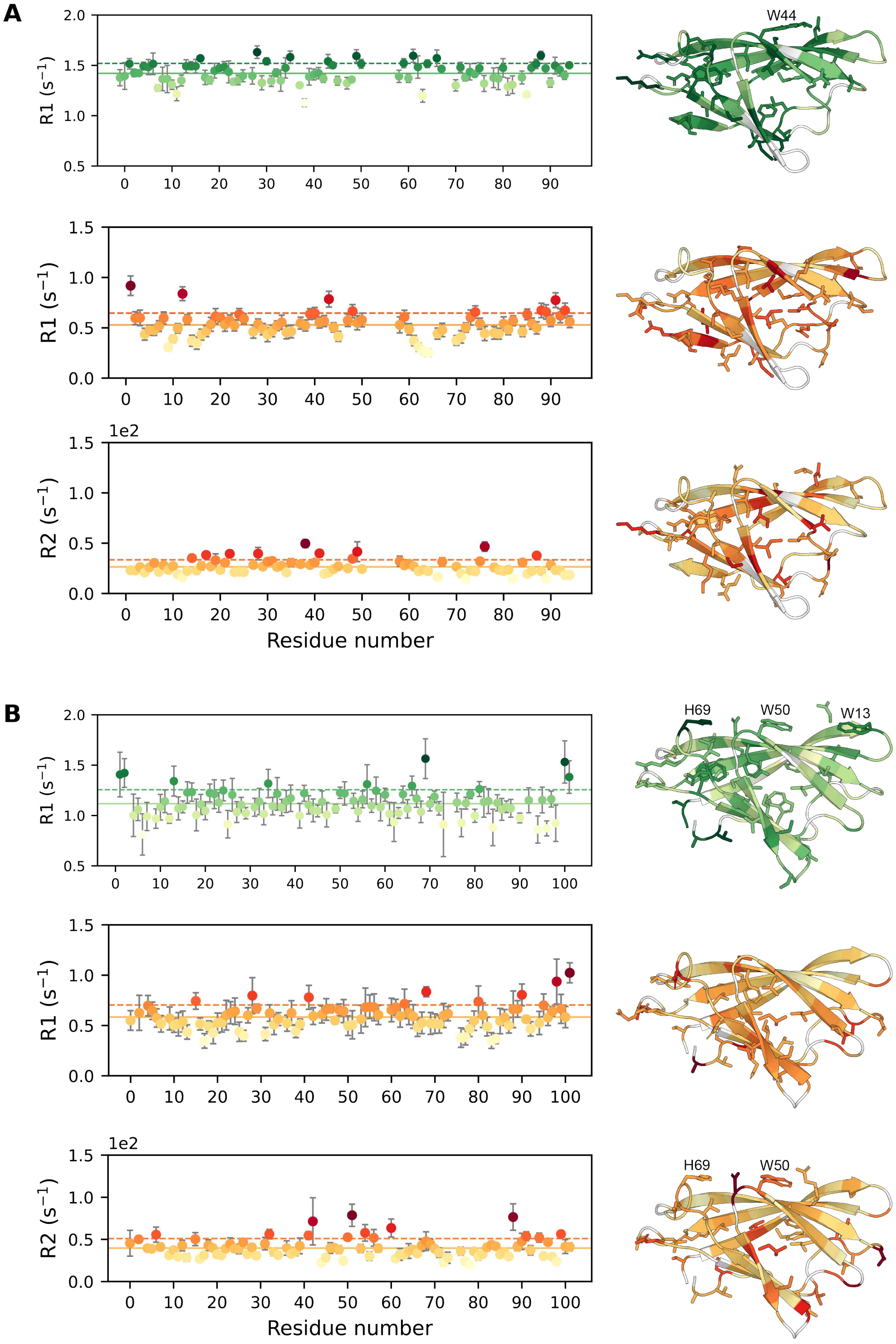


**Figure S14.** Nuclear spin relaxation rates measured for backbone resonances of *Ba*CBM2 (A) and *Sc*CBM2 (B). Upper panel: $^{15}$N $R_1$ relaxation, middle panel: $^{1}$H $R_1$ relaxation and lower panel: $^{1}$H $R_2$ relaxation. Solid lines show the average relaxation rate, and dotted lines represent the relaxation rate one standard deviation above the average. Relaxation rate magnitude is indicated by color gradients and shown on the respective AlphaFold structures on the right. Residues with relaxation rates above the average value are shown as sticks.

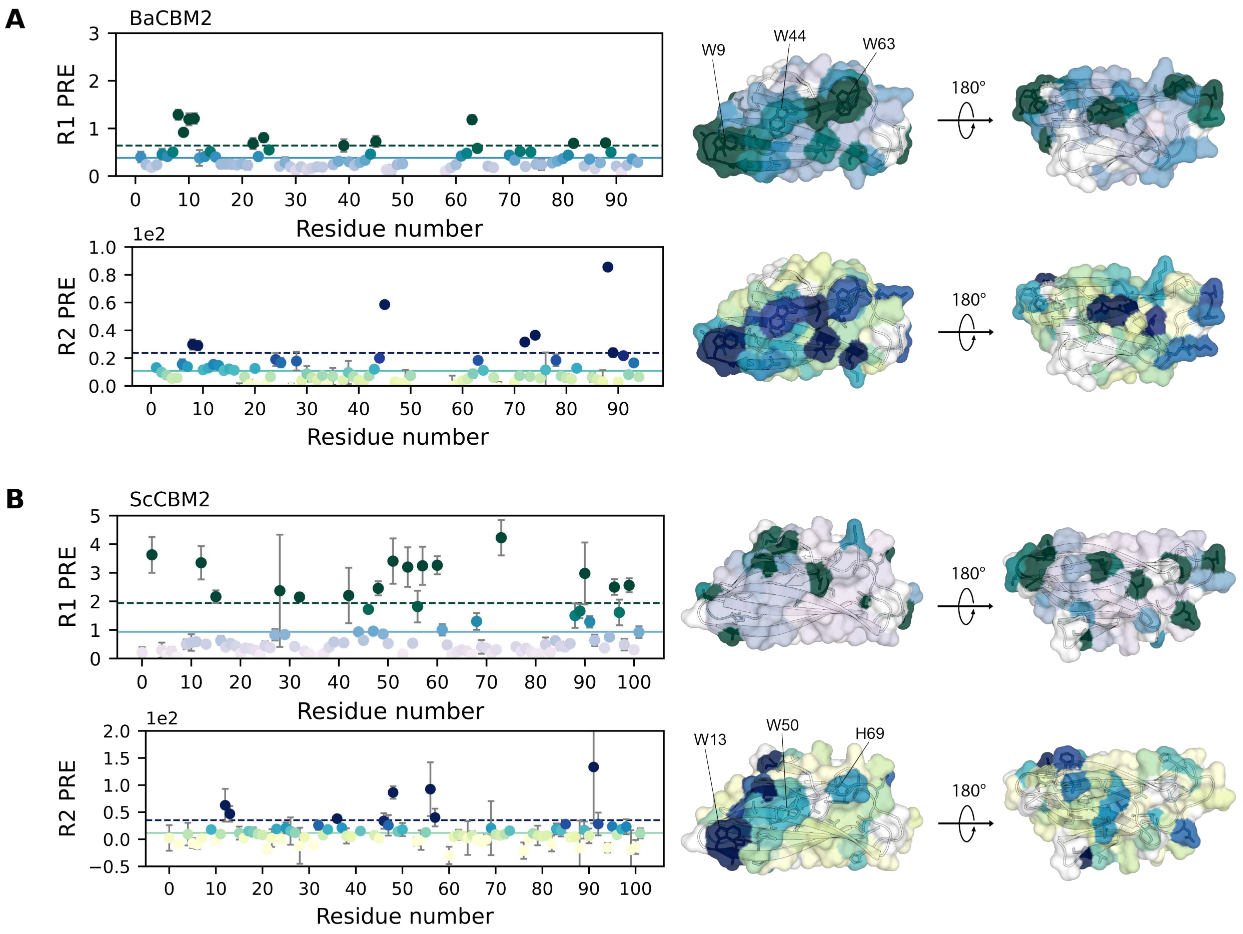


**Figure S15.** Solvent paramagnetic relaxation enhancement (sPRE) from gadodiamide in bulk solvent, measured for backbone resonances of *Ba*CBM2 (A) and *Sc*CBM2 (B). Upper panel: $^1$H $R_1$ sPRE and lower panel: $^1$H $R_2$ sPRE. Solid lines show the average sPRE, and dotted lines represent the sPRE one standard deviation above the average. sPRE magnitude is indicated by color gradients and shown on the respective AlphaFold structures on the right. Residues with sPRE above the average value are shown as sticks.

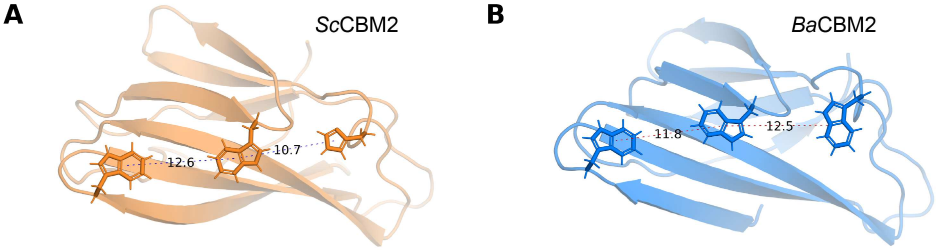


**Figure S16.** Simulated structures of *Sc*CBM2 (A) and *Ba*CBM2 (B), showing the distances between the rings of the aromatic triad.

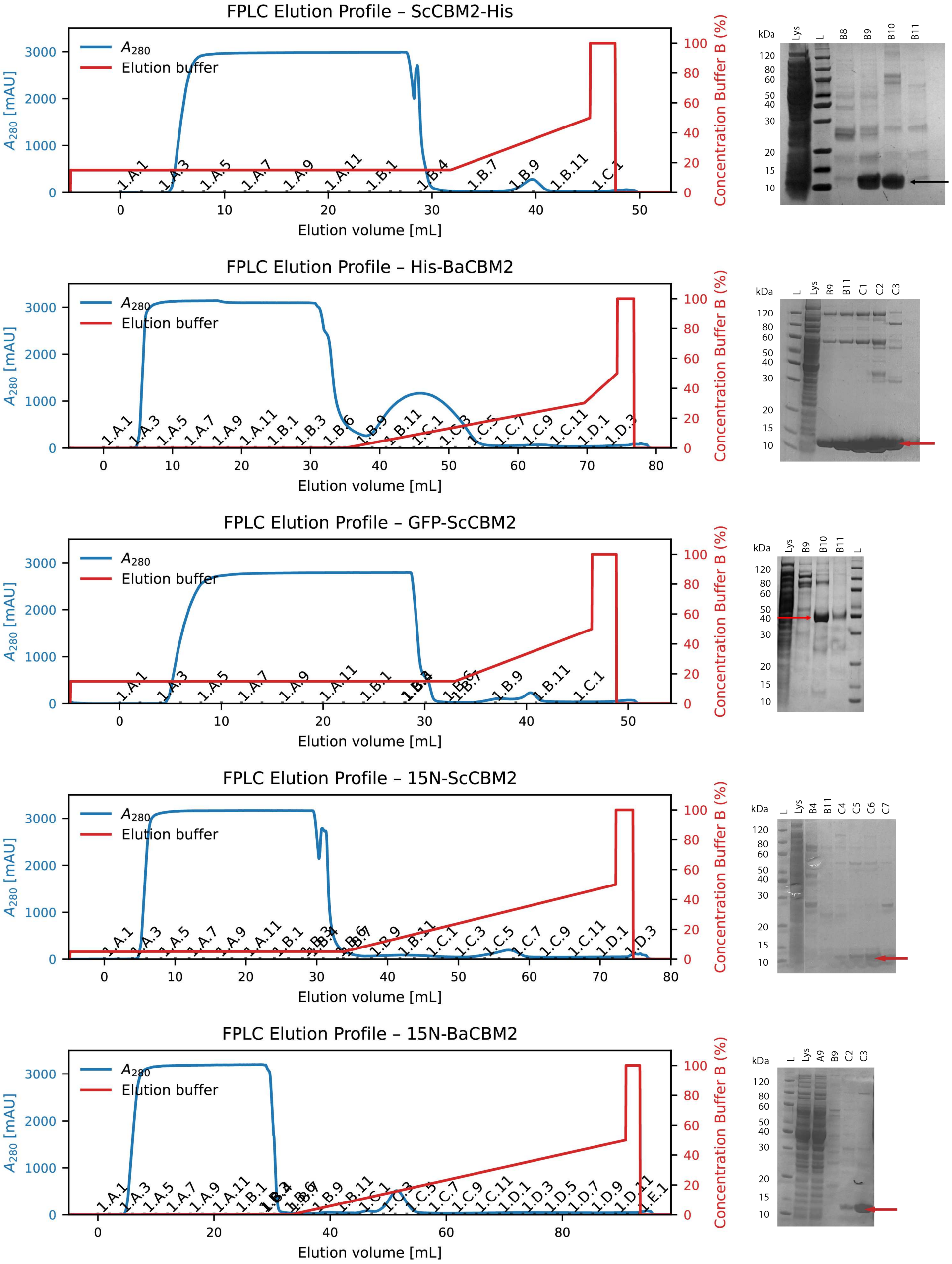


**Figure S17.** Chromatograms from Ni affinity FPLC purification and images of SDS-PAGE gels.

### References


[1] AP Rennison, et al., A high-throughput investigation of the binding specificity of carbohydrate-binding modules for synthetic and natural polymers. *ChemSusChem* **18**, e202500468 (2025).

[2] F Madeira, et al., The EMBL-EBI Job Dispatcher sequence analysis tools framework in 2024. *Nucleic Acids Res* **52**, W521–W525 (2024).

[3] GE Crooks, G Hon, JM Chandonia, SE Brenner, WebLogo: A sequence logo generator. *Genome Research* **14**, 1188–1190 (2004).

[4] J Abramson, et al., Accurate structure prediction of biomolecular interactions with AlphaFold 3. *Nature* **630**, 493–500 (2024).

[5] The PyMOL Molecular Graphics System, Version 3.1.5.1 (2025).

[6] J Weber, et al., Interaction of carbohydrate-binding modules with poly(ethylene terephthalate). *Applied Microbiology and Biotechnology* **103**, 4801–4812 (2019).

[7] Y Nishiyama, P Langan, H Chanzy, Crystal structure and hydrogen-bonding system in cellulose Iβ from synchrotron X-ray and neutron fiber diffraction. *Journal of the American Chemical Society* **124**, 9074–9082 (2002).

[8] JA Kaduk, PDF Card 00-056-1717. *Private communication* (2005).
